# Dynamic actin-mediated nano-scale clustering of CD44 regulates its meso-scale organization at the plasma membrane

**DOI:** 10.1101/751834

**Authors:** Parijat Sil, Nicolas Mateos, Sangeeta Nath, Sonja Buschow, Carlo Manzo, Kenichi G. N. Suzuki, Takahiro Fujiwara, Akihiro Kusumi, Maria F. Garcia-Parajo, Satyajit Mayor

**Affiliations:** National Centre for Biological Sciences, Bangalore, India; ICFO-Institut de Ciencies Fotoniques, The Barcelona Institute of Science and Technology, Barcelona, Spain; Institute of Stem cell and Regenerative medicine, Bangalore, India; Facultat de Ciències i Tecnologia, Universitat de Vic - Universitat Central de Catalunya, 08500 Vic, Spain; Centre for Highly Advanced Integration of Nano and Life Sciences (G-CHAIN), Gifu University, Japan; Institute for Integrated Cell-Material Sciences (WPI-iCeMS), Kyoto University, Japan; ICREA, Pg. Lluís Companys 23, 08010, Barcelona, Spain; Okinawa Institute of Science and technology, Graduate University, Japan; Manipal Institute of Regenerative Medicine, Manipal Academy of Higher Education, Bangalore, India; University of Utrecht, Netherlands

**Keywords:** actomyosin, cartography, CD44, fluorescence emission anisotropy, formin, homo-FRET, meshwork, meso-scale organization, nanoclustering, nano-scale organization, plasma membrane, single particle tracking

## Abstract

Transmembrane adhesion receptors at the cell surface, such as CD44, are often equipped with modules to interact with the extracellular-matrix(ECM) and the intra-cellular cytoskeletal machinery. CD44 has been recently shown to compartmentalize the membrane into domains by acting as membrane pickets, facilitating the function of signaling receptors. While spatial organization and diffusion studies of membrane proteins are usually conducted separately, here we combine observations of organization and diffusion by using high spatio-temporal resolution imaging on living cells to reveal a hierarchical organization of CD44. CD44 is present in a meso-scale meshwork pattern where it exhibits enhanced confinement and is enriched in nano-clusters of CD44 along its boundaries. This nanoclustering is orchestrated by the underlying cortical actin dynamics. Interaction with actin is mediated by specific segments of the intracellular-domain(ICD). This influences the organization of the protein at the nano-scale, generating a selective requirement for formin over Arp2/3-based actin-nucleation machinery. The extracellular-domain(ECD) and its interaction with elements of ECM do not influence the meso-scale organization, but may serve to reposition the meshwork with respect to the ECM. Taken together, our results capture the hierarchical nature of CD44 organization at the cell surface, with active cytoskeleton-templated nano-clusters localized to a meso-scale meshwork pattern.

## Introduction

Heterogeneity in the distribution of membrane proteins and lipids is becoming an increasingly appreciated paradigm in the context of the organization of molecules at the plasma membrane (Sezgin et al., 2017). This regulated, non-random distribution of membrane proteins such as signaling receptors, is implicated in their molecular function and signaling output (Garcia-parajo et al., 2014). The advent of super-resolution microscopy and breakthroughs in single molecule techniques has revolutionized our understanding of cellular organization at the molecular level (Klotzsch et al., 2013; Kusumi et al., 2005; van Zanten & Mayor, 2016). The major goal from such techniques has traditionally been to obtain detailed descriptions of protein clustering, cluster sizes or inter-molecular distances. However, these super-resolution techniques are often technically demanding and associated invasive sample preparation methods are fraught with criticism for being non-physiological. Additionally, although such studies of membrane constituents inform us on the organizational details at the molecular level, there have been fewer efforts to understand the organization and dynamics of proteins at larger spatial scales, to ascertain whether there exists any spatial hierarchy in membrane protein organization.

Studies of the membrane organization of many transmembrane receptors such as TCRs, EGFR, E-Cadherin, GPCRs or chemokine receptors such as CXCR-4, have advanced our understanding of changes at the nano-scale due to receptor dimerization or oligomerization (∼2-40 nanometers) in the presence or absence of the cognate ligand (Beck-garcía et al., 2015; Cohnen et al., 2016; Hofman et al., 2010; Martinez-Munoz et al., 2018; Overton & Blumer, 2000; Strale et al., 2015; Terrillon & Bouvier, 2004). At the same time, studies elucidating the inhomogeneous diffusion behavior of membrane proteins such as transferrin receptors (Kusumi and Sako, 1996) or CD44 (Freeman *et al*., 2018) have revealed the presence of compartments in the cell membrane at a larger length scale (∼ few hundred nanometers), templated by the underlying cytoskeletal meshwork. The potential hierarchy in the nature of organization of membrane proteins has been speculated in the past based on evidences from clustering and diffusion studies of different proteins (Kusumi *et al*., 2011). It is likely that a unified study of diffusion and organization interrogating the distribution of a particular membrane protein at different spatial scales will provide information of any underlying hierarchy in spatial scales of organization.

Type-1 trans-membrane proteins are a major and abundant class of integral membrane proteins that span three distinct environments, the extracellular space, transmembrane and cytoplasmic milieu. The lymphocyte homing receptor CD44, is a type I trans-membrane protein involved in cell-matrix adhesion (Ponta et al., 2003). It has a heavily glycosylated extracellular domain (ECD) that ensures binding to extra-cellular lectins such as galectins, besides being able to bind to its ligand hyaluronic acid (HA) as well as other components of the extra cellular matrix such as fibronectin and osteopontin (Ponta *et al*., 2003; Senbanjo and Chellaiah, 2017). Previous studies have shown that the ECD of CD44 is clustered by Galectin-3 which in turn also binds glycosphingolipids and is important for the endocytosis of the protein by a clathrin-independent pathway (Howes *et al*., 2010; Lakshminarayan *et al*., 2014). Additionally, HA binding has been shown to influence the dynamics of the protein at the plasma membrane (Lakshminarayan *et al*., 2014; Freeman *et al*., 2018). The juxta-membrane O-glycosylation site and the trans-membrane region with two putative palmitoylation sites confer the ability on the protein to partition into detergent resistant membrane fractions or cholesterol enriched domains on the plasma membrane (Thankamony and Knudson, 2006; Shao *et al*., 2015).

At the intracellular side, the relatively short 70 amino acids long cytoplasmic tail of CD44 interacts with multiple cytoskeletal adaptor proteins. The association of the protein with ezrin has been shown to be important for T cell migration in interstitial spaces of endothelial cells (Mrass *et al*., 2008). The interaction with ezrin also influences the protein’s ability to act as membrane picket in macrophages providing a functional partitioning of the FcγRIIA at the plasma membrane and facilitating its phagocytic function in macrophages (Freeman *et al*., 2018). Ankyrin binding has been shown to be important for HA binding by CD44 (Bourguignon, 2008). A proteomic analysis of the interacting partners of the CD44 cytoplasmic tail has also revealed an interaction with other cytoskeletal adaptors such as vinnexin, IQGAP1 and talin1 (Skandalis et al., 2010). The modularity of these potential cytoskeletal interactions in the tail of CD44 via its multiple cytoskeletal adaptor binding sites opens up possibilities to study how they may independently regulate organization and turn-over of the protein at the cell surface.

Thus, the diverse structural attributes of CD44 impart this receptor with the ability to be influenced by extracellular interactions, membrane composition and the actin cytoskeleton. Hence, it also provides an ideal platform to uncover general principles of how such molecules are organized at varying length scales, determined by distinct modes of interaction in the different milieu and also the interplay between these length scales. Nevertheless, studies so far have not systematically investigated the role of the different structural domains of the protein in the organization and dynamics of the liganded as well as the native un-liganded receptor on the membrane, at multiple spatial scales.

In this study we have exploited various imaging methods in living cells to characterize the organization of CD44 at the single molecule level over multiple spatiotemporal scales. Single molecule tracking at different labeling densities allowed us to capture the dynamics of CD44, both at the nano- and meso-scale levels. We define nano-scale organization as being built of individual molecules brought together within ∼10nm scale and meso-scale as domains ∼ 100 nm - <1µm in scale. By means of interleaved homo-FRET-based anisotropy and high-density single molecule imaging, we show that the meso-scale organization of CD44 is significantly associated with its nanoclusters. Moreover, homo-FRET anisotropy measurements revealed a role for the actomyosin machinery and formin, which is also reflected in the mesoscale organization. Overall, our data provide evidence for a hierarchical organization of CD44, wherein each layer of organization is determined by distinct interactions of the receptor.

## Results

### Spatio-temporal mapping of CD44 reveals a mesh like distribution of the protein at the mesoscale

To explore the dynamics of CD44 with high spatiotemporal resolution we utilized the standard isoform of mouse CD44 (Ponta *et al*., 2003) tagged with a SNAP domain at the N-terminus and GFP at the C-terminus (SNAP-CD44-GFP) (Fig 1a, b, Table 2). This chimeric protein can be labeled at the extra-cellular side using cell-impermeable benzylguanine (BG)-conjugated fluorophores that covalently link to the extracellular SNAP domain. SNAP-CD44-GFP was expressed in wild type mouse embryonic fibroblast (MEFs) cells that endogenously express CD44 as well as produces the ligand HA (Gerecht et al., 2007; Siiskonen et al., 2015) and labeled with SNAP-Alexa 546 (or BG-Alexa 546). Sub-saturation labeling conditions (≤30 nM) were required for performing single particle tracking (SPT) in order to unambiguously reconstruct all the individual receptor trajectories of diffusion. However, this approach under-samples the cell membrane and thus provides little information on membrane regions dynamically explored by the receptor (Fig. 1c). We thus increased the labeling density (∼50-100 nM), to ensure higher sampling frequencies of the membrane protein and yet maintaining the ability of detecting individual molecules in each single frame to determine their coordinates with sub-pixel accuracy (Fig. 1c’).

**Figure 1:**
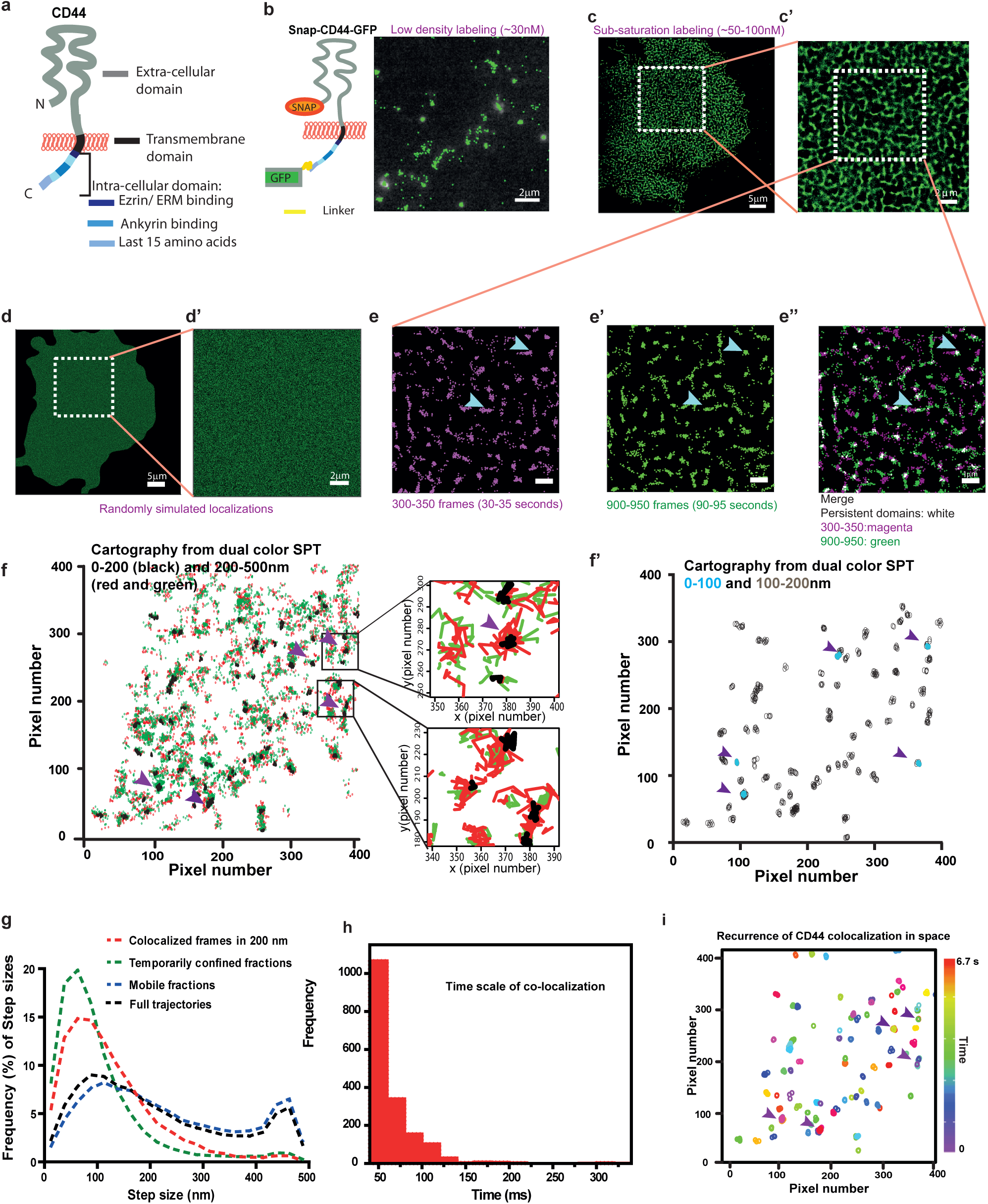
CD44 exhibits a non-random distribution at the plasma membrane at multiple spatiotemporal scales. (a) Schematic of a standard isoform of CD44 showing key domains of the protein, namely, the extra-cellular domain(ECD), transmembrane domain (Tm) and the intra-cellular domain(ICD). (b) Schematic of SNAP-CD44-GFP and representative dynamic cartography of CD44 obtained at sub-labeling conditions (∼30nM). Each dot corresponds to the (x,y) co-ordinates (with sub-pixel accuracy) of individual receptors as they diffuse on the cell membrane. The (x,y) coordinates over 50 sequential frames are collapsed and overlaid in to a single fluorescence frame. (c) Cartography of SNAP-CD44-GFP obtained at higher labeling conditions (∼50-100 nM); (x,y) co-ordinates from 1000 frames (1354066 localizations) collapsed in a single map with a zoomed-in ROI(c’). (d) Simulated cartography with similar number of localizations as in (c), distributed in a random fashion, and enlarged ROI. (e) Cartography construction of (x,y) coordinates in the marked ROI in (c) from 50 consecutive frames obtained at two different experimental time windows, between 30s-35s (magenta, e) and 90s-95s (green, é) and merged image (right, e”). Blue arrowheads highlight regions of confinement, and white dots represent persistent confinement regions or sites re-visited by the receptor. (f) Cartography obtained from 400 frames (∼6.7 s) of DC-SPT data obtained by co-labeling CD44 with JF549-cpSNAP ligand and JF646 SNAP ligand. Green and red dots correspond to localizations with the two different dyes indicating inter-particle distances between 200-500 nm and black dots correspond to inter-particle distances < 200nm. Zoomed-in ROIs depict the indicated reconstructed trajectories; f’ shows the subset corresponding to localization of particles with 0-200nm inter-particle distance (black in f) where the inter-particle distance corresponds to <100nm (blue circles) and 100-200nm (grey circles). Note the <100nm co-localization events always correspond to regions where localizations at the larger length scale of 100-200nm and 200-500 nm inter-particle distance are also found (indicated by purple arrowheads in f and f’). (g) Frequency distribution of step sizes of particles from trajectories wherein particles exhibit co-localization (red), temporary arrest (green); mobile (blue) and from the full trajectories (black) (27856 trajectories). (h) Frequency distribution of duration of co-localization of particles identified in Fig 1f. Note the lifetime of co-localization is in the range of < 100ms. (i) 2D plot of all co-localized particles (<200 nm) obtained from the trajectories identified in panel (f), where purple arrowheads indicate colocalization events which occur repeatedly at the same spot over the time period of observation. Color LUT bar indicates observation time from 0-6.7 seconds.

Time-lapse images were acquired at 10 frames per second (fps) for 1000 seconds and the spatial coordinates of identified individual molecules over multiple frames were collapsed into a single frame to obtain a time-dependent cartography of the regions dynamically explored by the receptor as described in an earlier study (Torreno-pina et al., 2014). This density regime offers the possibility of building up a large number of localizations to construct dynamic meso-scale cartography of CD44 distribution over the entire cell membrane (Fig 1c’). Remarkably, we found that CD44 diffusion and distribution is largely inhomogeneous, describing a clear mesh-like spatiotemporal distribution at the meso-scale (1c’, zoomed-in). This mesh is defined by regions frequently re-visited by the receptor and/or induced by its temporal arrest on the cell membrane. This is in stark contrast with the distribution of simulated randomized localizations on the plasma membrane (Fig. 1d, 1d’) which appears homogeneous at the same length scale. Indeed, enlarged regions of the cartography, from the same patch of the cell membrane, generated at two different time windows, show the dynamic character of the mesh (Fig. 1e, 1e’), and importantly, reveal sites of confinement/trapping of the receptor, evidenced by the large number of localizations (>10^6^ for Fig. 1c’) occurring within regions between ∼90-200 nm in size (Fig. S1b). Moreover, some of these regions have a long persistence time (∼50-60 seconds, Fig. 1e, 1e’ and merged image in 1e’), indicating that the receptors could be stably confined in these regions and/or transiently tether repeatedly to the same regions. Similar experiments conducted in cells which exhibit very low surface levels of endogenous CD44 (COS-7 cells (Fig. S1a, a’)) and the extracellular ligand, HA (Knudson *et al*., 1993; Shyjan *et al*., 1996) (CHO cells (Fig S1c), also yielded similar results. Together, these results indicate that CD44 is organized in a mesh-work pattern on the plasma membrane and this distribution is independent of binding to its ligand HA on the extra-cellular side or surface levels of endogenous proteins.

Since the experiments were conducted on the surface of the cell close to the coverslip, it is conceivable that the observed meso-scale pattern visualized for CD44 is an artifact of the patterning of the membrane due to its adhesion to the cell substrate. To rule this out, we imaged mouse embryonic fibroblasts (MEFs) expressing SNAP-CD59-GPI, a GPI anchored protein, unrelated to CD44. Analysis of the meso-scale map of SNAP-CD59-GPI also reveals a meso-scale meshwork pattern on the cell surface, indicating a compartmentalized state of the plasma membrane (Fig S1f). To additionally rule out the possibility that over-expression of the chimeric SNAP-tagged CD44 protein induces such a distribution, we investigated how endogenous CD44 is organized at the plasma membrane by labeling the protein using anti-CD44 antibody and performing Stochastic Optical Reconstruction Microscopy (STORM) in fixed cells (Fig S1d). Endogenous CD44 at non-adherent membrane of the lamella, away from the adhesion surface, also revealed a meshwork-like pattern of the protein at the meso-scale. STORM revealed a nano-scale clustered distribution of CD44 laid out in a non-random mesoscale mesh-like pattern at the cell membrane. Nearest neighbor distance analysis on CD44 clusters of multiple STORM images further confirmed that the nano-clusters of CD44 are distributed in a manner distinct from simulated randomized distribution of nano-clusters (Fig S1e). Therefore the meshwork like pattern of CD44 reflects a hitherto unappreciated intrinsic organization of this protein in the membrane of living cells.

In order to discriminate between single and/or multiple receptors being confined, as obtained in the cartography, we then turned to dual color single particle tracking (DC-SPT), by using sub-saturation labeling conditions (Kusumi et al., 2005). For this, we labeled the SNAP-CD44-GFP expressed in MEFs, using two different dyes (JF549-cpSNAP and JF646-SNAP ligands) and tracked the motion of the receptor at 60 fps for 400 frames (6.7 sec) (Supp. Video1 and 2). Localization maps created from superposing 400 frames of the DC-SPT images revealed typical trajectories of Brownian diffusion interspersed with transiently confined trajectories of the single color tracks (Fig. S2a). Analysis of >2500 trajectories reveal the existence of a large percentage of transiently confined receptor (68.4±2.3%) on the cell membrane with majority of confinement time restricted to ≤∼3seconds (Table 1 and Fig S2b). When we examined the DC-SPT data, we observed a noticeable overlap of CD44 molecules (co-localized) that reside in confined regions (Fig. 1f: purple arrowheads indicate black dots in the map, and corresponding trajectories). To quantify specific co-localization, we determined the occurrence of co-localized events as a function of inter-particle distances within defined areas (depicted as radius on the X axis in Fig S2c), and compared the results to those of diffusion from randomized trajectories (obtained from 180 degrees flipped images of the same regions). A random distribution is expected to have an inter-particle distance distribution index equal to 1, with values greater and smaller indicating clustering and dispersion respectively (Clark and Evans, 1954). From the inter-particle distance quantification, we defined co-localized particles, as those which exhibited inter-particle distances (between two differently labeled SNAP-CD44-GFP molecules) less than 200 nm for 3 consecutive frames. We also observe a subset of these events to correspond to inter-particle distance <100nm (a length scale more precisely matched with combined localization precision of the fluorophores) (Fig 1f’). Interestingly, the length scale over which CD44 exhibits co-localization corresponds to the length scale over which it exhibits transient confinement as is evident from analyzing the step size distribution (Fig. 1g). We also quantified the co-localization lifetime and find that individual co-localization event lasts for <100ms (Fig 1h). Temporal analysis of localization events revealed recurrence of co-localization events at the same spatial co-ordinates over a period of 400 frames (0-6.7 seconds) (Fig. 1i, color indicating time at which co-localization occurred), indicating hotspots of trapping of same/different pairs of receptors. These data thus indicate the existence of hotspots on the plasma-membrane that can both restrict the diffusion of CD44 and recruit multiple CD44 molecules. Moreover, the cartography analysis, STORM and DC-SPT data (Fig. S1d and S1e, 1f) suggest the formation of CD44 clusters that might be organized in a meso-scale meshwork on the plasma membrane.

**Table 1:** CD44 diffusion characteristics from single particle tracking

| Protein Chimera | Percentage transient confinement | Trap radius in nanometers(nm) | Diffusion co-efficient in $\mu\text{m}^2/\text{sec}$ |
| --- | --- | --- | --- |
| SNAP -CD44 -GFP | $68.4 \pm 2.3$ | $42.5 \pm 1.6$ | $0.02 \pm 0.01$ (slow:~(69±4)%)<br>$0.16 \pm 0.01$ (fast: ~(31±10)%) |
| SNAP -CD44TmICD-GFP | $57.1 \pm 4.2$ | $54.9 \pm 4.6$ | $0.02 \pm 0.01$ (slow:49.3±3.2)%)<br>$0.18 \pm 0.004$ (fast: 50.7±9.9)%) |
| SNAP-CD44Tm-GFP | $45.4 \pm 8.3$ | $68.9 \pm 11.7$ | $0.06 \pm 0.02$ (slow: 29.5±9.5)%)<br>$0.23 \pm 0.01$ (fast: 70.5±4.4)%) |

**Table 2:**
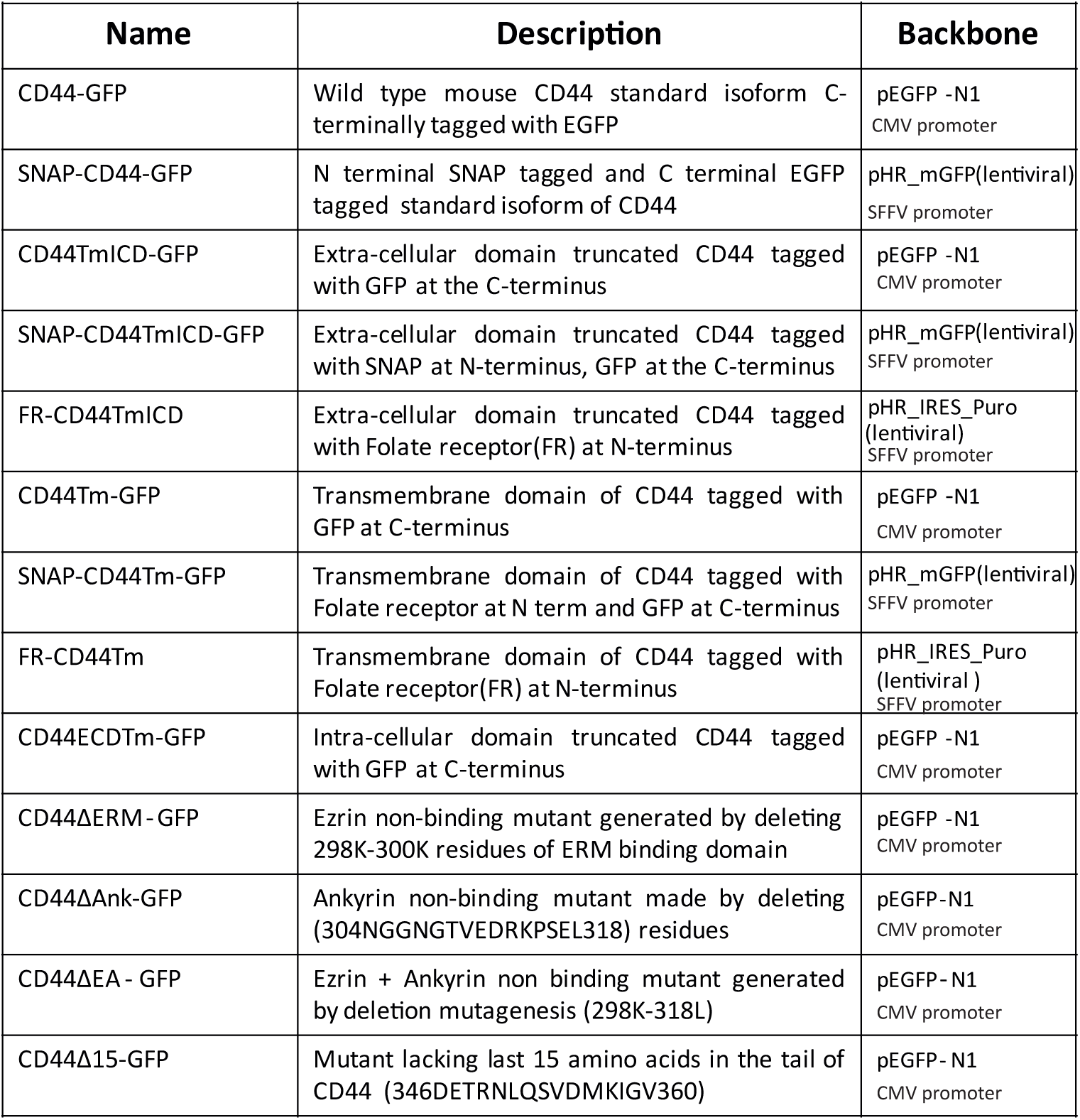
List of different constructs used in the study

### The dynamic meso-scale meshwork of CD44 is composed of nanoclusters

STORM imaging of endogenous CD44 in CHO cells as described above, as well as an earlier study (Lakshminarayan *et al*., 2014), provide evidence for the existence of nanoscale clusters of CD44 at the plasma membrane. Moreover, CD44 exhibits a high incidence of co-localizations (<200nm) in DC-SPT as well as spatially confined localizations that emerge as a mesh-like pattern in the cartography analysis. Together, these observations motivated us to investigate the clustering interactions of CD44 at the nano-scale using homo FRET microscopy (Ghosh et al., 2012).

Fluorescence emission anisotropy based homo-FRET measurements probes the proximity of fluorescently-tagged proteins at a molecular length scale ∼ Forster’s radius, (∼5 nm for the GFP fluorophore (Ghosh *et al*., 2012)) on the living cell membrane, reporting molecular interactions at a length scale ∼10 times smaller than achievable resolution in STORM. Using this method we identified regions of low and high anisotropy in the membrane of un-perturbed living cells in four different cell types: COS-7 cells (Fig. 2a, S3d, S3d’), CHO cells (Fig. 3b, 3d), MEFs (Fig. S3f, f’) and MCF-7 (Fig S3e, e’), each of which has different properties. While CHO and MEFs express endogenous CD44, COS-7 and MCF-7 cells have very low surface levels of endogenous protein (Fig S3g), and both COS-7 and CHO cells do not synthesize a major ECM component, HA, that can bind CD44 from the extracellular side (Shyjan *et al*., 1996; Yang *et al*., 2012). The regions of low anisotropy correspond to an enrichment of CD44-GFP molecules at ≤ 5 nm inter-molecular distances, thus indicating the occurrence of nanometer scale encounters of CD44 molecules on the cell membrane at a steady state. These results corroborate the co-localization observed by DC-SPT as well as spatially confined localizations observed in the cartography.

**Figure 2:**
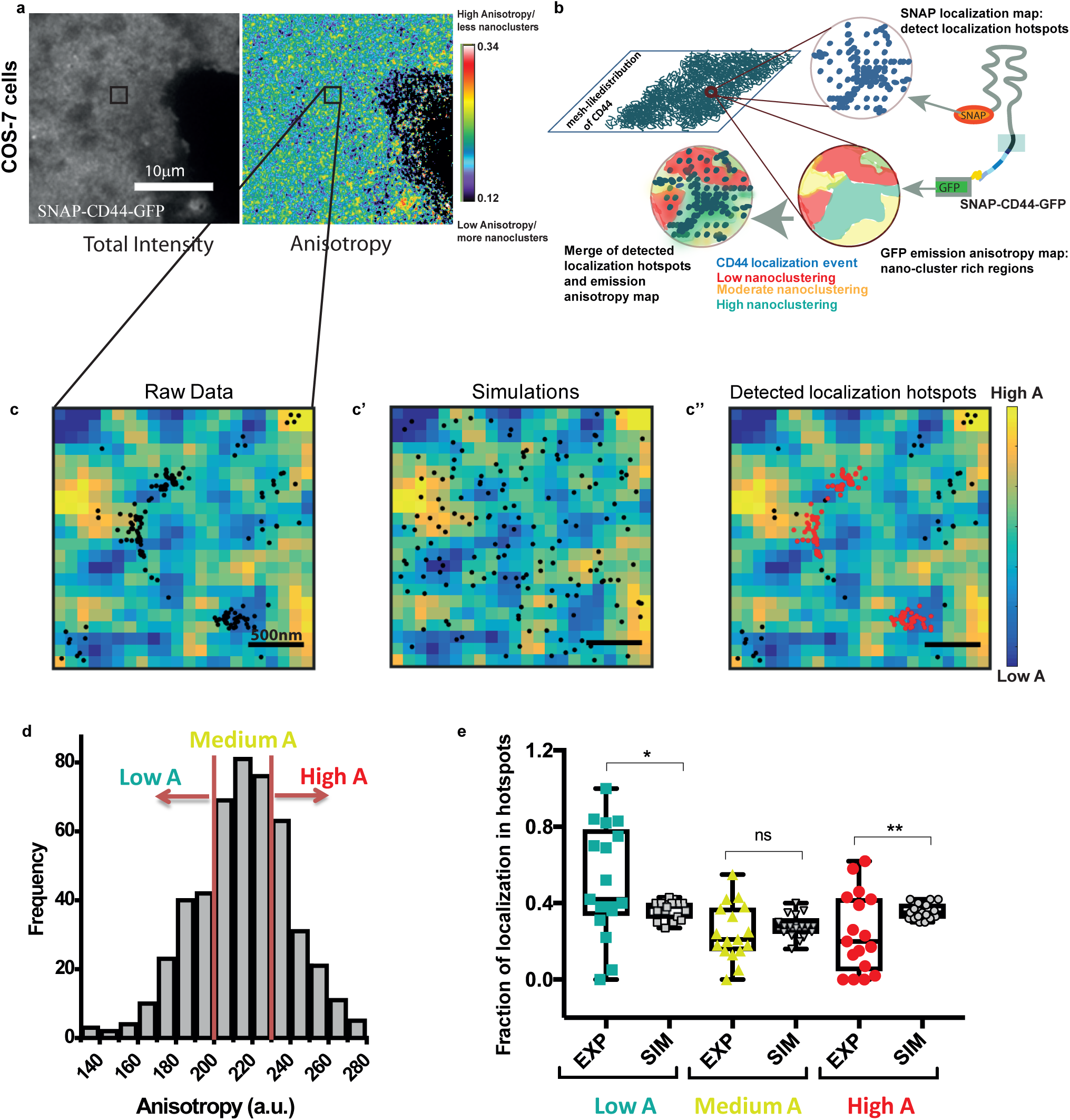
Meso-scale meshwork of CD44 co-localizes with regions enriched in CD44 nanoclusters. (a) Total GFP fluorescence intensity and anisotropy map of the SNAP-CD44-GFP protein expressed in COS-7 cells that exhibit low levels of surface CD44. Note that the anisotropy image shows regions of low anisotropy (blue) and high anisotropy (red), corresponding respectively, to regions enriched in, or depleted of CD44 molecules in nanoscale proximity (CD44 nanoclusters). (b) Schematic depicting the methodology by which FRET based anisotropy maps was correlated to localization maps obtained from high density single molecule imaging and cartography analysis. (c, c’, c”) Representative ROI image depicting the anisotropy map overlaid with localizations from raw cartography images integrated over 40 frames (left), random localizations obtained from simulations (center) and detected localization hotspots (red dots) of SNAP-CD44-GFP (right). (d) Histogram of the anisotropy values for the ROI shown in panel (c). Red vertical lines indicate the thresholds chosen to classify regions of low anisotropy (Low A), medium anisotropy (Medium A) and high anisotropy (High A), where medium anisotropy is binned around the median value of anisotropy in a given ROI. (e) Fraction of detected localizations in the ‘localization hotpots’ in low, medium and high anisotropy regions compared to simulated localizations. Each symbol in the plot corresponds to a single ROI, and the data is obtained from at least 6 different cells from>15 ROIs. Difference between distributions has been tested using Kolmogorov-Smirnov test.

**Figure 3:**
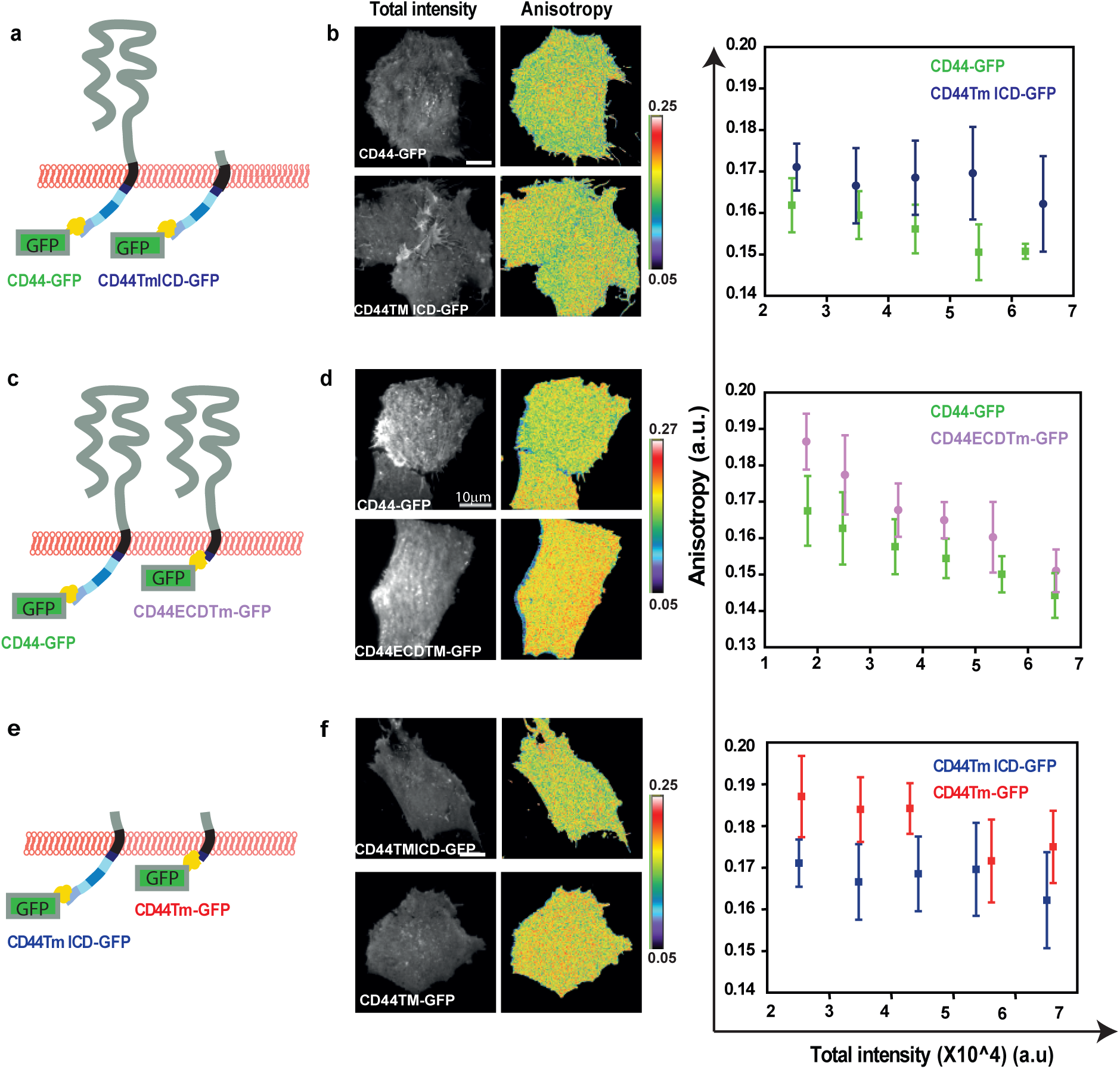
ECD and ICD independently affect CD44 nanoclustering. Schematics (a,c and e) depict CD44-GFP constructs expressed in CHO cells, used to generate the corresponding intensity and anisotropy images in (b, d and f). Anisotropy versus intensity plots show a significant increase in anisotropy in the truncated protein lacking the extra-cellular domain (a, b; p= < 10-43), intracellular domain (c, d; p<10-58), or when the construct lacking the extra-cellular domain (data from the same experiment as a and b) is compared to one lacking both extracellular domain and the intracellular domain (e, f; e<10-77). All raw distributions are statistically significant by Mann-Whitney test for each condition. (The data is from one representative experiment. Fig. 3b: CD44-GFP= 20 fields, CD44TmICD-GFP = 27 fields; Fig. 3d: CD44-GFP = 25 fields, CD44ECDTm-GFP = 13 fields; Fig. 3f: CD44Tm-GFP = 15 fields).

To ascertain the relationship between nano and meso-scale dynamic organization of CD44, we expressed the SNAP-CD44-GFP construct in COS-7 cells to obtain fluorescence emission anisotropy maps from the GFP tag on the SNAP-CD44-GFP, interleaved with single molecule imaging data from the sub-saturation labeled SNAP tag, amenable for generating cartography. We chose COS-7 cells since they exhibit low levels of CD44 at the cell surface and also upon ensuring that these cells exhibit nano-clustering of ectopically expressed CD44-GFP (Fig S3d, S3d’ and S3g) (Jiang *et al*., 2002; Yang *et al*., 2012). We selected different anisotropy ROIs and super-imposed the corresponding spatial coordinates of individual molecules integrated over 40 frames (20 frames preceding and 20 frames following the anisotropy image) (Fig. 2b, 2c). We restricted our analysis to windows of 40 frames around an anisotropy image to reduce temporal variations that might occur between the anisotropy and cartography (see Methods). We then identified spatially restricted enriched localizations, termed „localization hotspots’ on the cartography maps and classified these „localization hotspots’ according to the corresponding anisotropy value (see experimental Methods and Fig. 2c, c’. c’’, 2d). A significantly higher fraction of localization hotspots were localized to regions of low anisotropy and correspondingly such localization hotspots were consistently depleted from the high anisotropy regions when compared to randomly dispersed localizations (Fig. 2e). These data indicate that the meso-scale regions observed on the cartography overlaps with the regions of increased nanoscale clustering of the receptor. As a whole, our results reveal a multi scale organization of CD44 on the cell membrane with the distribution of nanoscale clusters correlated to the meso-scale meshwork. This motivated an exploration of the mechanism behind the formation of the nano-clusters of CD44.

### The extra- and intra-cellular domains of CD44 independently affect nanoclustering of CD44 at the plasma membrane

To probe the mechanism(s) responsible for the organization of CD44 molecules at nanoscale proximity, we examined both intensity dependence and spatial anisotropy distribution of various mutants of CD44-GFP (Fig. 3a, c, e; Table 2) for the description of the different constructs used) expressed in HA deficient CHO cells, by fluorescence emission anisotropy based homo-FRET microscopy. Fluorescence emission anisotropy of CD44-GFP was intensity dependent indicating a concentration dependent change potentially due to: (a) protein-protein interactions, (b) potential dilution by endogenous CD44, (c) combination of both (Fig. 3b). The later possibility was confirmed by using MCF-7 cells which have very low levels of cell surface CD44, where fluorescence emission anisotropy of CD44GFP exhibited visibly lower intensity dependence while at the high intensity range, it became concentration dependent (Fig S3e, e’). These observations suggest that at the lower expression range of CD44-GFP in cells with significant endogenous CD44, the intensity dependence of its anisotropy is a convolution of both, dilution by endogenous unlabelled protein as well as concentration-dependent protein-protein interactions. However, at higher levels of expression, protein-protein interactions and trivial density dependent FRET may contribute to the intensity dependence of anisotropy. Consistent with this, deletion of the ECD of CD44 (CD44TmICD-GFP) resulted in an increase in anisotropy and reduced its intensity dependence (Fig. 3a, b), consistent with an attenuation of concentration dependent interactions as compared to the full length receptor in CHO cells. While these cells do not synthesize HA (Shyjan *et al*., 1996), CD44 expressed on the surface of these cells can still bind galectins (Lakshminarayan *et al*., 2014) and may have other protein-protein interactions mediated by the ECD of the receptor. These interactions could lead to a concentration-dependent clustering, which is reduced by deletion of the ECD. Thus, the prominent intensity dependence and lower fluorescence emission anisotropy exhibited by the full length receptor as compared to the mutant likely results from ECD interactions of CD44, impacting its nano-scale organization.

To ascertain if the deletion of the ECD completely abolished CD44 nano-clustering, we measured the change in anisotropy of the fluorescently labeled CD44TmICD protein upon dilution of fluorophores by photo-bleaching. Since enhanced GFP is capable of reversible photo-bleaching, giving rise to artifacts in bleaching-based homo-FRET measurement (Sinnecker et al., 2005), we resorted to a different strategy for labeling the truncated CD44 with a fluorophore exhibits reduction in energy transfer efficiency upon destruction of FRET competent fluorophores by bleaching (Sharma *et al*., 2004). We designed a chimeric Folate Receptor (FR)-tagged version of the ECD truncated protein (Fig. S3a). This chimeric construct was expressed in CHO cells and labeled with a fluorescently labeled folate analog (PLB^TMR^: N^α^-pteroyl-Nɛ-Bodipy^TMR^-L-lysine) (Goswami *et al*., 2008), and then imaged while photo-bleaching the labeled cells. If the labeled proteins are clustered, the emission anisotropy of FR-CD44TmICD should increase since photo-bleaching reduces the concentration of fluorescent proteins engaged in energy transfer (Sharma *et al*., 2004). PLB^TMR^-labelled FR-CD44TmICD exhibited an increase in emission anisotropy upon photo-bleaching (Fig. S3b, c), indicating that the ECD truncated protein retains the ability to engage in nanometer scale homomeric interactions at the plasma membrane. The slope in the anisotropy plot is an indication of the extent of nanoclustering, i.e., higher the slope, the greater is the extent of nanoclustering (Sharma *et al*., 2004). Overall these results indicate an inherent ability of CD44TmICD to nanocluster on the cell membrane, and the extent of clustering in CD44 is also modulated by interactions in the extracellular milieu.

The findings described above led us to investigate the role of the ICD in CD44 nanoclustering. For this, we measured the fluorescence emission anisotropy of the full-length receptor (CD44-GFP) and a CD44 mutant lacking only the ICD or cytoplasmic tail (CD44ECDTm-GFP) (Fig. 3c). The results indicated that the full-length wild type protein is clustered to a greater extent compared to the ICD truncated protein, as indicated by the lower anisotropy values obtained with the full-length protein (Fig. 3d). The truncated protein still retains a concentration-dependent anisotropy, consistent with the possibility of passive interactions affecting its nano-clustering. Similar increase in anisotropy values were obtained in COS-7 and MCF-7 cells transfected with the same constructs (Fig. S3e, e’), indicating that the results obtained in the CHO cells (S3d, d’), were minimally affected by the endogenous, unlabeled CD44 population at the cell surface. Consistent results were also obtained in MEF cells that secrete HA and express significant levels of endogenous CD44, indicating that the disruption of nano-clustering due to loss of the ICD in these cells is strong enough to manifest as significant increase in anisotropy, in-spite of the presence of the polymeric ligand HA in the extra-cellular milieu as well as potential fluorophore dilution due to co-clustering of labeled CD44 with endogenous unlabelled CD44 proteins (Fig S3f, f’).

To further validate the clustering potential of the cytoplasmic domain, we deleted the entire ICD in the CD44TmICD-GFP construct to create a transmembrane domain only protein (CD44Tm-GFP) (Fig. 3e). We found that the anisotropy of the resultant protein increased compared to the CD44TmICD-GFP (Fig. 3f), consistent with the clustering potential of the ICD. Differences in nanoclustering in the presence and absence of the ICD were further corroborated by comparative photo-bleaching analysis of folate receptor (FR) tagged FR-CD44TmICD and the truncated FR-CD44Tm in CHO cells. We found that FR-CD44TmICD is clustered to a greater extent than FR-CD44Tm (Fig. S3a, b, c), as indicated by a reduction in the slope of the „anisotropy vs. normalized intensity’ curve of the trans-membrane FR-CD44Tm as compared to the FR-CD44TmICD protein.

The results show that ECD and ICD independently affect CD44 nanoclustering. The ECD has a greater impact in establishing passive interactions with partners on the cell membrane giving rise to a strong intensity/ expression level dependent clustering of CD44 at the cell surface. Even though, the transmembrane region appears to have small but detectable ability to nanocluster the receptor (due to a minor residual slope in the photo-bleaching analysis), it is the ICD that strongly enhances the nano-clustering ability of CD44.

### CD44 nanoclustering correlates with its tethering strength on the plasma membrane

To further understand how CD44 nanoclustering affects the lateral diffusion of the receptor we carried out SPT at sub-saturation labeling conditions (∼30nM) on the full length SNAP-CD44-GFP (Supp. Video 3 and 4), and the truncated SNAP-CD44TmICD-GFP (Supp. Video 5) and SNAP-CD44Tm-GFP (Supp. Video 6) constructs in MEFs cells (Fig. 4a). These cells are also ideally suited for testing the effect of the extracellular influence of HA which may affect CD44 dynamics at the membrane. Individual trajectories for the three different constructs were obtained (Fig. 4b, Fig S4a) and the fraction of mobile trajectories were quantified (Fig. 4c; calculated from escape probability of molecules in MEFs; Fig. S4c, Fig. S4d are in COS-7 cells; Table 1); trajectories with diffusion coefficients < 0.02 m^2^/s were defined as immobile. Deletion of the ECD increased the fraction of mobile receptors as compared to the full length protein (Fig. 4c), an effect that became even more pronounced with further removal of the ICD. Moreover, analysis of the transient confinement areas showed tighter regions of confinement for the SNAP-CD44-GFP and SNAP-CD44TmICD-GFP as compared to the SNAP-CD44Tm-GFP mutant (Fig. 4d, Fig. S4e and Table 1) and the overall diffusion coefficients were significantly slower for the full length receptor (Fig. 4e, Table1). The results indicate that interactions by the ECD ensure slower diffusion and the cytoplasmic domain ensures both slower diffusion as well as tighter confinement of CD44, at the plasma membrane. The difference between the wild type protein and mutant lacking ECD is more pronounced in MEFs compared to COS-7 cells, potentially owing to the presence of HA in the matrix of MEFs, consistent with the observations made in the earlier study by Freeman et al (2018).

**Figure 4:**
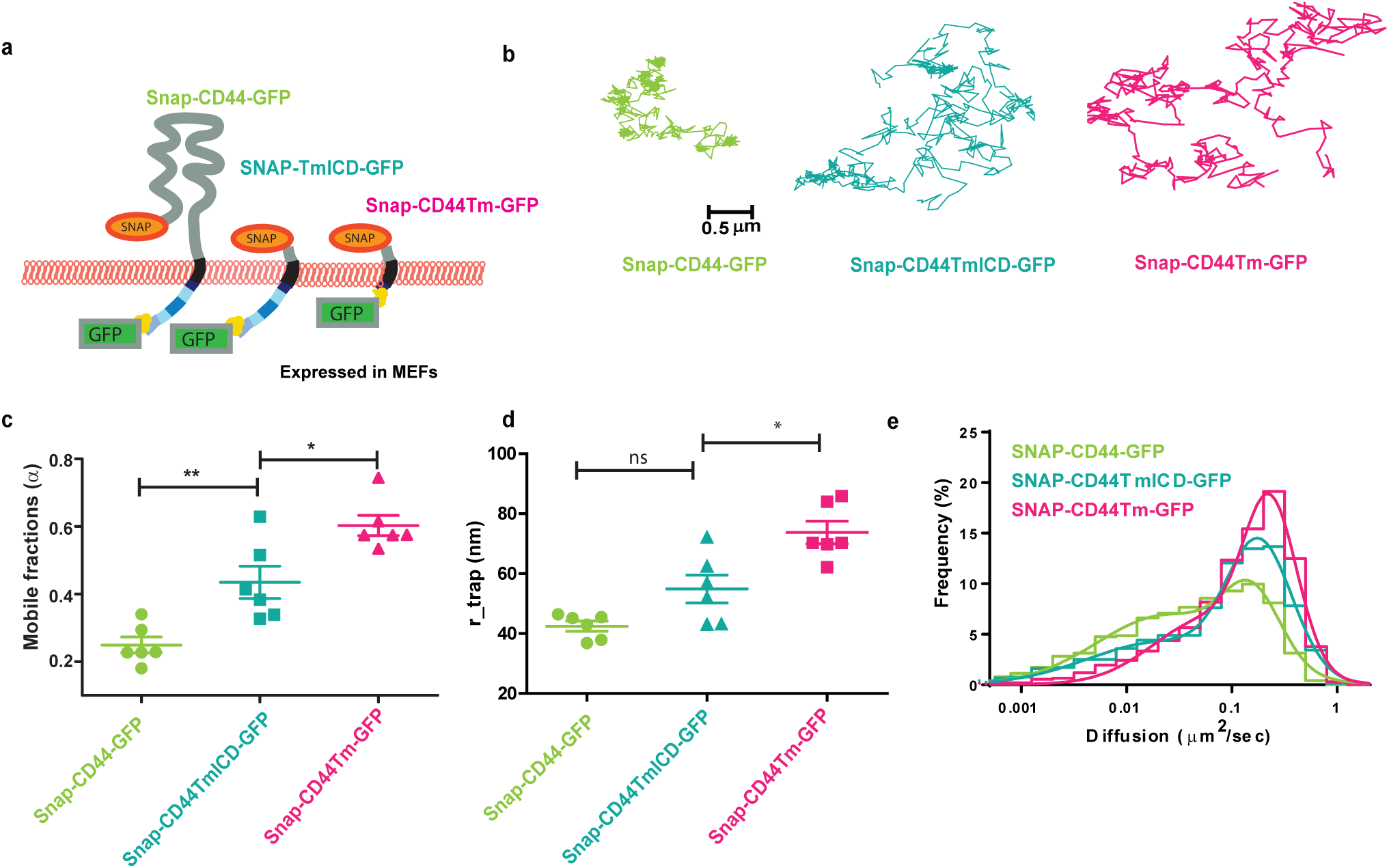
Extent of CD44 nanoclustering correlates with the strength of tethering on the cell membrane. (a) Schematic show SNAP-tagged constructs expressed in MEFs, utilized for SPT. (b) Representative trajectories for the indicated constructs show distinct diffusion characteristics of the different constructs; (c-e) Quantification of the (c) mobile fraction by escape probability method, (d) confinement radius (r_trap), (e) and diffusion coeffi cients of the full length and the truncated mutants. The data is derived from at least 6 cells for each construct. Number of trajectories: SNAP-CD44-GFP= 2977; SNAP-CD44TmICD-GFP = 2783; SNAP-CD44Tm-GFP = 4744.

To further elucidate the consequences of the differences in tethering strength of the wild type and the transmembrane mutant as observed in the single color SPT experiments, analysis of co-localization events and quantification of inter-particle distance using DC-SPT, on both the full length receptor and the transmembrane mutant lacking both the ECD and ICD, further corroborated that interactions by these domains can affect the co-localization propensity of the protein (Fig. S2e, S2e’, S2f, S2f’). Together with the anisotropy data (Fig. 3, Fig. S3) these results point to a strong correlation between the degree of CD44 nanoclustering and its tethering at the cell membrane: the full length receptor exhibits the strongest nanoclustering (as derived from the fluorescence anisotropy analysis) and tighter confinement and/or tethering at the cell membrane. On the other hand, deletion of both the ECD and cytoplasmic tail reduces nanoclustering and increases the mobility of the receptor, with reduced tethering at the membrane (Table 1).

### Meso-scale organization of CD44 is influenced by its cytoplasmic interactions

Since CD44 nanoclustering is spatially correlated to its meso-scale distribution, we then tested whether alteration in the nanoclustering potential of the different mutants correlates with the manifestation of any defects in their meso-scale organization. SNAP-CD44-GFP, SNAP-CD44TmICD-GFP and SNAP-CD44Tm-GFP constructs were expressed in MEFs, exogenously labeled and imaged at a temporal resolution of 10 fps as described earlier, in order to generate cartography of the different constructs (Fig. 5a). Visual inspection of the cartography already show more tightly bound localizations in the case of the full length receptor and a larger number of dispersed localizations for the SNAP-CD44Tm-GFP mutant. Comparison of the confinement areas revealed similar confinement strength for the full length receptor (0.028±0.013) µm^2^and the mutant lacking the ECD (0.027±0.013) µm^2^ (Fig. 5b, c), indicating that the ECD does not play a major role on the meso-scale organization of the receptor. Consistent with these results, we did not find significant differences on the fractional number of localizations found on the meshwork between the full length receptor (SNAP-CD44-GFP) and the mutant lacking the ECD(SNAP-CD44TmICD-GFP) (Fig. 5d). In contrast, the mutant lacking the cytoplasmic tail as well as the ECD (SNAP-CD44Tm-GFP) exhibited larger confinement areas (0.032±0.013) µm^2^ (Fig. 5b,c) and a significantly lower number of localizations associated to the meshwork as compared to the full length receptor (SNAP-CD44-GFP) or the mutant lacking the ECD alone (SNAP-CD44TmICD-GFP) (Fig. 5d). This result strengthens the observation from SPT, that, the cytoplasmic domain mediates tight confinement of the receptor at the plasma membrane.

**Figure 5:**
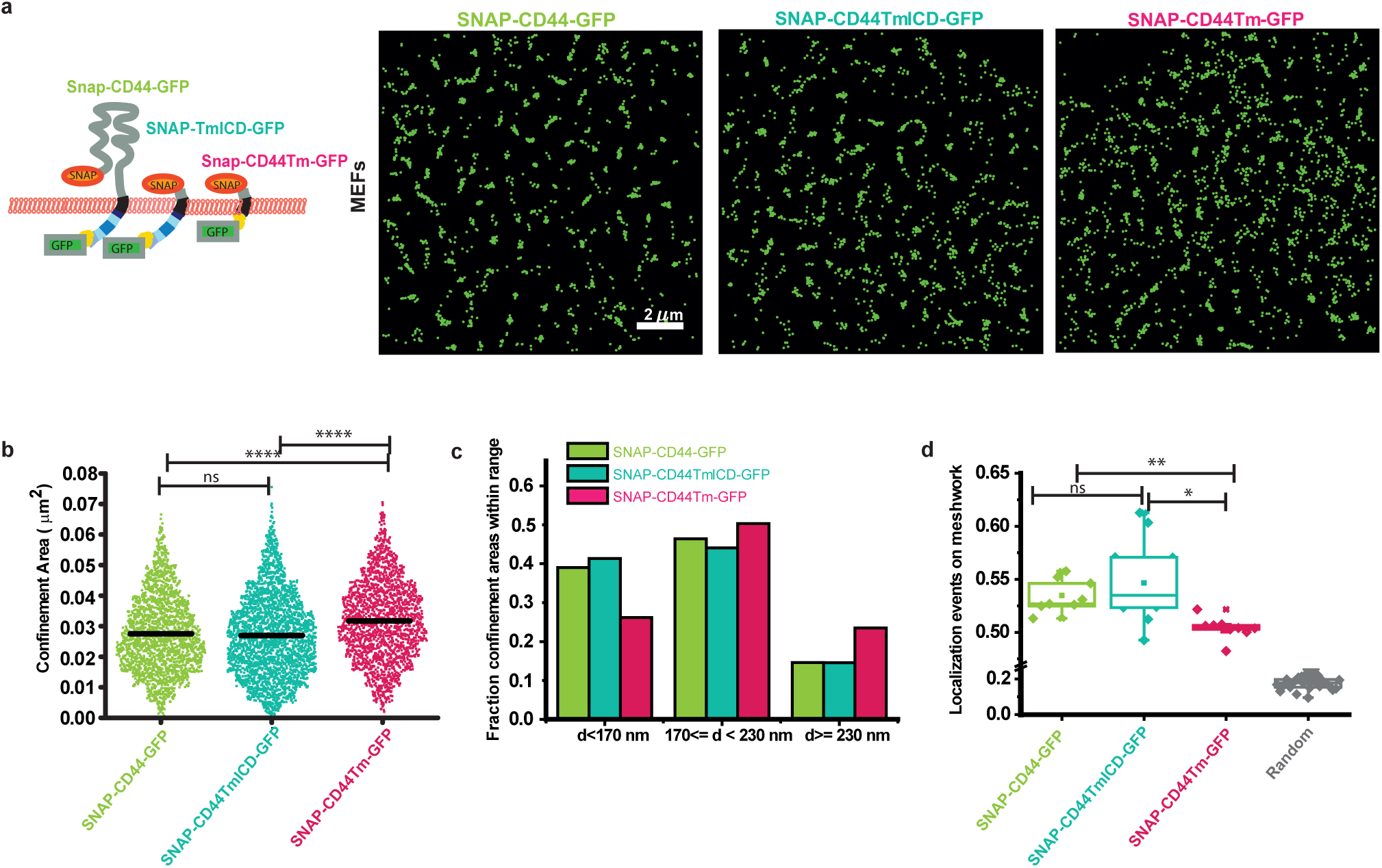
Meso-scale organization of CD44 is determined primarily by interactions of the intra-cellular domain. (a) Representative cartography maps of the indicated CD44 constructs expressed in MEFs obtained from imaging at 10fps and accumulating the spatial coordinates of individual molecules over 2s (20 frames). (b) Quantification of the confinement areas for the different constructs during 2s. Black lines correspond to the mean value. (c) Relative fractions of confinement areas for the different constructs, classified as a function of the confinement length, i.e., d<170nm, 170<d<230nm or d>=230nm. (d) Fraction of localization events that belong to the meshwork for the different constructs and compared to the fraction of similar type of localizations measured from randomized localizations. The data is from one representative experiment. The experiment has been conducted at least twice with similar results. Data were obtained from a number of cells expressing SNAP-CD44-GFP (8), SNAP-CD44TmICD-GFP (11) or SNAP-CD44Tm-GFP (9). Difference between distributions was tested for significance using Kruskal-Wallis and post hoc test with Tukey-Kramer. (b): SNAP-CD44-GFP & SNAP-CD44TmICD-GFP: p=0.258 (ns); SNAP-CD44-GFP & SNAP-CD44Tm-GFP: p <e-9; SNAP-CD44TmICD-GFP & SNAP-CD44Tm-GFP: p<e-9. (d): SNAP-CD44-GFP & SNAP-CD44TmICD-GFP: p=0.8564 (ns); SNAP-CD44-GFP & SNAP-CD44Tm-GFP: p < 0.005; SNAP-CD44TmICD-GFP & SNAP-CD44Tm-GFP: p=0.0218. SNAP-CD44Tm-GFP(n) = 9 cells, SNAP-CD44TmICD-GFP (n) = 11 cells, SNAP-CD44-GFP (n) = 8 cells.

We also performed similar experiments in HA deficient COS-7 cells and obtained comparable results (Fig. S5). Since the confinement areas and number of localizations associated to the meshwork result from multiple re-visiting and/or arrest of the receptor to the underlying meshwork, these results strongly suggest that the mutant lacking both the ECD and the cytoplasmic tail (SNAP-CD44Tm-GFP) compared to the mutant lacking the ECD alone (SNAP-CD44TmICD-GFP), is less tethered to the meshwork. Of note, we also performed simulations of random localizations and overlaid them to an experimentally obtained meshwork to obtain a “basal” fraction of localizations that are stochastically found over the meshwork (labeled as random in Fig. 5d). Comparison with the *in-silico* generated data revealed that even in the absence of the cytoplasmic tail, the SNAP-CD44Tm-GFP mobility is somewhat constrained by this underlying mesh albeit to a lower extent than the cytoplasmic domain containing counterparts. Therefore, our results strengthen the arguments for cytoplasmic interactions as a major player in orchestrating the nano-and meso-scale organization of CD44. Since the cytoplasmic tail of CD44 interacts with multiple cytoskeletal adaptor proteins such as ezrin and ankyrin (Bourguignon, 2008; Mrass *et al*., 2008), our results, suggest that CD44 nanoclustering might be induced by its tethering to the actin cytoskeleton. This finding resonates with the recently published results of CD44 in macrophages where diffusion characteristics of the protein are affected by tethering to the cytoskeleton mediated by ezrin (Freeman *et al*., 2018) and leads us to investigate the role of the actin cytoskeleton in the nano-clustering as well as the meso-scale organization of the protein.

### Nanoclustering of CD44 is regulated by actin dynamics

Previous work has shown that actin binding confers the ability of proteins to associate with the actomyosin-clustering machinery in living cells. Here, dynamic actin filaments driven by myosin, propel the formation of actin asters, driving the generation of clusters of proteins that associate with these structures (Gowrishankar *et al*., 2012). Since CD44 has been shown to engage with the cytoskeleton by binding to ezrin and ankyrin via its cytoplasmic tail (Bourguignon, 2008; Mori *et al*., 2008; Donatello *et al*., 2012), we investigated whether actomyosin perturbations would affect the clustering of the receptor. Here we investigated the effects of actomyosin perturbations in CHO cells, since the ICD of CD44 was found to support nano-clustering of CD44 in all the cell types tested. Firstly, we treated CHO cells with the actin filament stabilizer Jasplankinolide (Jas) to create blebs which represent membranes devoid of the dynamic actin cortex (Jaumouillé *et al*., 2014). Fluorescence emission anisotropy of CD44-GFP on blebs of Jas-treated cells was higher compared to the flat membranes of untreated cells (Fig. 6a). This also holds true for the CD44TmICD-GFP mutant, which is devoid of the extra-cellular domain, (Fig. S6a). These observations strongly suggest that the interactions with a dynamic actin cortex (absent in blebs), is a key determinant of nano-clustering of the protein at the cell surface. Moreover, treatment of cells with a cocktail of inhibitors (ML-7 and Y27632/ H11152) (Totsukawa *et al*., 2000; Saha *et al*., 2015) that inhibit myosin regulatory light chain phosphorylation of class II non-muscle myosins, thereby inactivating them, resulted in a loss of nano-clustering of the CD44-GFP, as indicated by the increase in emission anisotropy of CD44-GFP compared to control cells (Fig. 6b and Fig. S6c: where similar results are also obtained for the ECD deleted mutant CD44TmICD-GFP). This result indicates that a dynamic actomyosin-driven mechanism facilitates nanoclustering of CD44 at the plasma membrane.

**Figure 6:**
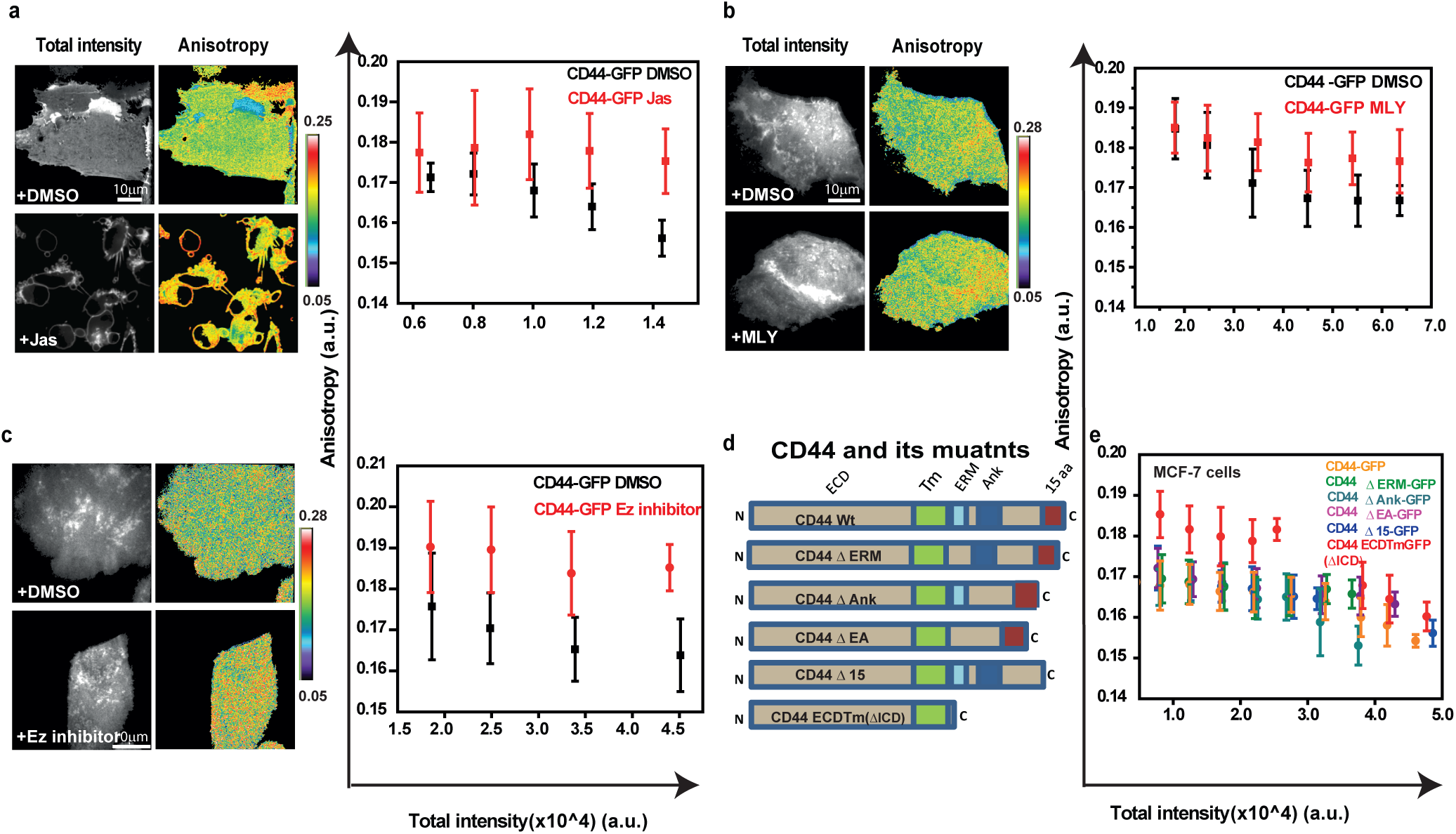
CD44 nanoclustering is regulated by the underlying actomyosin machinery. Total intensity and anisotropy images of cells expressing CD44-GFP (a-c) expressed in CHO cells, either untreated or treated with actin polymerization stabilizer, Jasplakinolide (a; Jas; 14 µM, 15 min; Con (n) = 10 fields, Treatment (n) = 22 fields), Myosin inhibition cocktail (b; MLY 20µM; 60 min; Con (n) = 20 fields, Treatment (n) = 26 fields), Ezrin inhibitor (c; 25µM; 60 min, Con (n) = 16 fields, Treatment (n) = 11 fields),. Graphs show anisotropy values plotted against intensity collected from regions from the cells as detailed in experimental methods. In all conditions treatment with the indicated inhibitors show a significant difference in the recorded values of anisotropy (p< 10-5), Difference between distributions has been tested for significance by Mann-Whitney tests. The data is from one representative experiment. Each experiment was conducted at least twice with similar results. (d) Schematic of CD44 and different deletion mutants for ezrin, ankyrin and last 15 amino acids of the tail with the names of the constructs indicated next to its diagram. (e) Plot shows intensity versus anisotropy distributions of the CD44 mutants in MCF-7 cells which exhibit low surface levels of CD44. (Distribution of anisotropy values were tested for significance using Mann-Whitney test and p<10-120 was obtained for CD44-GFP and CD44ECDTm-GFP; CD44-GFP (n) = 19 fields, CD44-ECDTm-GFP (n) = 15 fields, CD44-Δ15GFP (n) = 16 fields, CD44-ΔERM-GFP(n) = 13 fields, CD44-ΔEA-GFP (n) = 16 fields, CD44-ΔAnk-GFP (n) = 17 fields).

CD44 has been shown to associate with the actin cytoskeleton binding proteins, ezrin and ankyrin (Bourguignon, 2008; Mrass *et al*., 2008) while the last 15 amino acids of CD44 confers it the ability to interact with talin1, vinnexin, LMO1 and IQGAP1, all of which are potential interactors of the actin cytoskeleton as well as multiple other proteins (Skandalis *et al*., 2010). To understand whether CD44 is associated with any particular adaptor protein that confers it with a cytoskeleton-sensitive clustering, we used two strategies. One where we perturbed the cytoskeletal coupling of CD44 using a small molecule inhibitor of ezrin function, and the other using site directed mutagenesis to specifically generate mutants that would be deficient in one or more actin binding domain: CD44ΔERM (deletion of ezrin binding site), CD44ΔAnk (deletion of ankyrin binding site), CD44ΔEA (both the ezrin and ankyrin binding sites are deleted) and CD44Δ15 (deletion of the last 15 amino acids), tagged to GFP on the cytoplasmic side (see Table 2; Fig. 6d).

When we inhibit ezrin function using the small molecule inhibitor of ezrin (NSC668394), it resulted in an increase of the fluorescence emission anisotropy of CD44 (Fig 6c), indicating the importance of ezrin function in CD44 nanoclustering. Similar effects were also observed for the CD44TmICD-GFP mutant (Fig. S6b). However, when we expressed the various truncation mutants in cells, homo-FRET based anisotropy measurements revealed minimal difference in steady state anisotropy distribution between the full length receptor and the mutant proteins in MEFs, CHO cells (Fig. S7a, S7a’, respectively), and validated in COS-7 and MCF-7 cells (Fig. S7a’’ and Fig. 6e) to ensure that smaller differences in the nano-clustering of the mutants compared to the wild type protein were also detected. This suggests that there are redundant ways of the mutant protein to associate with the actin-myosin machinery, and it is only when the entire cytoplasmic tail is deleted that this engagement is lost and nanoclustering abrogated.

### Meso-scale organization and turnover of CD44 is regulated by formin-nucleated actin dynamics

The diffusion of CD44 has been suggested to be sensitive to formin-generated actin filaments (Freeman *et al*., 2018) since upregulation of Rho activity (which in turn regulates formin activity), influences the diffusion behavior of CD44. In order to test which actin nucleation machinery is responsible for CD44 nanoclustering, we inhibited formin and Arp2/3 mediated actin filament-nucleation activity in CHO cells using small molecule inhibitors, SMI-FH2 and CK-666 respectively. CD44 nanoclustering was much more sensitive to inhibition of formin nucleation (Fig 7a) compared to Arp2/3 perturbation (Fig S6d). These results indicate that formin-nucleated F-actin filaments not only influence the mobility of the receptor as reported previously (Freeman et al 2018), but importantly, also promotes its nanoclustering, and as a consequence may also influence its meso-scale organization.

**Figure 7:**
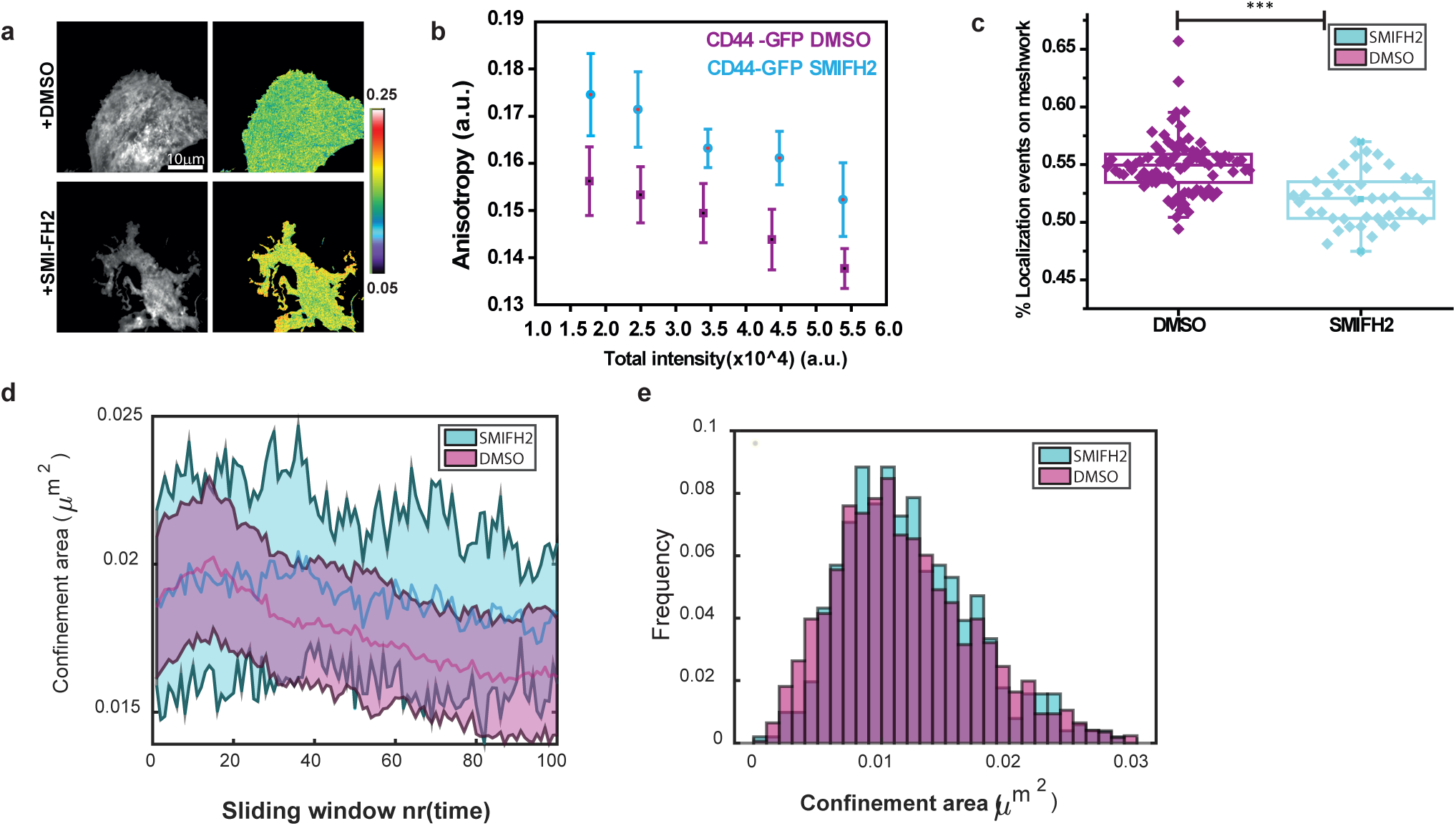
Formin mediated Actin polymerization affect nano as well as meso-scale distribution and turnover of CD44. (a, b) Total intensity and anisotropy images of cells expressing CD44-GFP expressed in CHO cells treated with formin inhibitor (SMIFH2 10µM, 30 min; Con (n) = 19 fields, Treatment (n) = 13 fields, p<10-5). (c) Plot describing fraction of localizations detected on the meshwork in control cells compared to formin inhibited condition (p<e-8) (d) Plot depicting time evolution of meso-sale domains upon vehicle (DMSO) versus formin inhibitor treatment. X axis depicts time as 2 seconds sliding window (depicted as frame number) and Y axis depicts confinement area. (e) Plot depicting confinement area of the mesoscale domains in formin perturbed cells compared to untreated ones do not exhibit detectable differences. (DMSO(n)= 12 cells, SMIFH2(n) = 9 cells).

To ascertain the effect of formin perturbation on the meso-scale meshwork we conducted high density single particle imaging of SNAP-CD44-GFP, as described before, in COS-7 cells where we earlier elucidated the co-existence of nanoclusters with meso-scale domains. Our results indicate that meso-scale meshwork of CD44 is perturbed in formin perturbed cells. Although the confinement area distribution is not significantly altered in formin perturbed cells compared to DMSO treated cells (Fig 7e), the fraction of localization events detected along the meshwork in formin-treated cells (Fig 7b, 7c, 7d) is significantly reduced which is reminiscent of the distribution of the SNAP-CD44Tm-GFP that lacks both the cytoplasmic and exoplasmic domains, and is also defective in nanoclustering.

A striking difference in the formin-treated cells compared to the untreated was in the turnover time of the meso-scale domains. Time evolution analysis of the meso-scale domains revealed that while untreated (vehicle treated) cells exhibited a visible disassembly/reorganization of the mesoscale domains, formin-treated cells exhibited a marked persistence of meso-scale domains (Fig 7d) during the observed time window. These results indicated that dynamic remodeling of the meso-scale meshwork is dependent on formin activity, consistent with the suggestion that formin driven actin polymerization is a key contributor to dynamic remodeling of the actin meshwork (Fritzsche *et al*., 2013).

## Discussion

CD44 has a multitude of extra-cellular and cytoplasmic interactions that makes it an ideal candidate for studying regulation of the organization of a typical membrane protein. Here we have used non-invasive methods to study nanoclustering and dynamics of CD44 using live-cell compatible techniques such as homo-FRET imaging and SPT methods to generate spatial maps of the protein at the plasma membrane at the nano and meso-scale. Previous studies have attempted to understand CD44 organization by multiple approaches, from characterizing graded distribution of GP-80 in motile fibroblasts (Ishihara et al., 1988) to super-resolution imaging wherein CD44 was found clustered at the cell membrane using STORM, and extracellular galectins were found responsible for their nanoclustering (Lakshminarayan *et al*., 2014). In another study, the ICD was implicated in supporting mobile clusters at the membrane based on hetero-FRET measurements, brightness number analysis, and biochemical cross-linking studies in mammalian cells (Wang *et al*., 2014). In a more recent study, SPT on CD44 revealed that CD44 diffusion is confined to pickets and fences, and may indeed determine the corralling of other membrane proteins such as the FcγRIIA (Freeman *et al*., 2018).

The results reported here provide a comprehensive understanding of the organization of CD44 by combining the determination of distribution and diffusion behavior of the protein across varying spatial scales at the plasma membrane of living cells. Cartography analysis (to probe the meso-scale organization of the protein) and its correlation with anisotropy measurements (reporting on nanoclustering), for the first time, bridges the gap between SPT based diffusion studies and the steady-state nanocluster detection method of homo-FRET. Complemented with the cartography analysis of single particle localizations and nanocluster distribution in STORM images, the combination of these approaches enabled us to build a hierarchical framework for the organization of a type-1 transmembrane protein at the plasma membrane (Fig. 8). We find that actomyosin templated nano-clusters of CD44 spatially enrich the receptors along a meso-scopic meshwork pattern, laid down by frequent localizations of the protein at the plasma membrane. These nanoclusters resemble actomyosin-based clusters observed for model transmembrane proteins with actin-binding domains (Chaudhuri et al., 2011; Gowrishankar et al., 2012).

**Figure 8:**
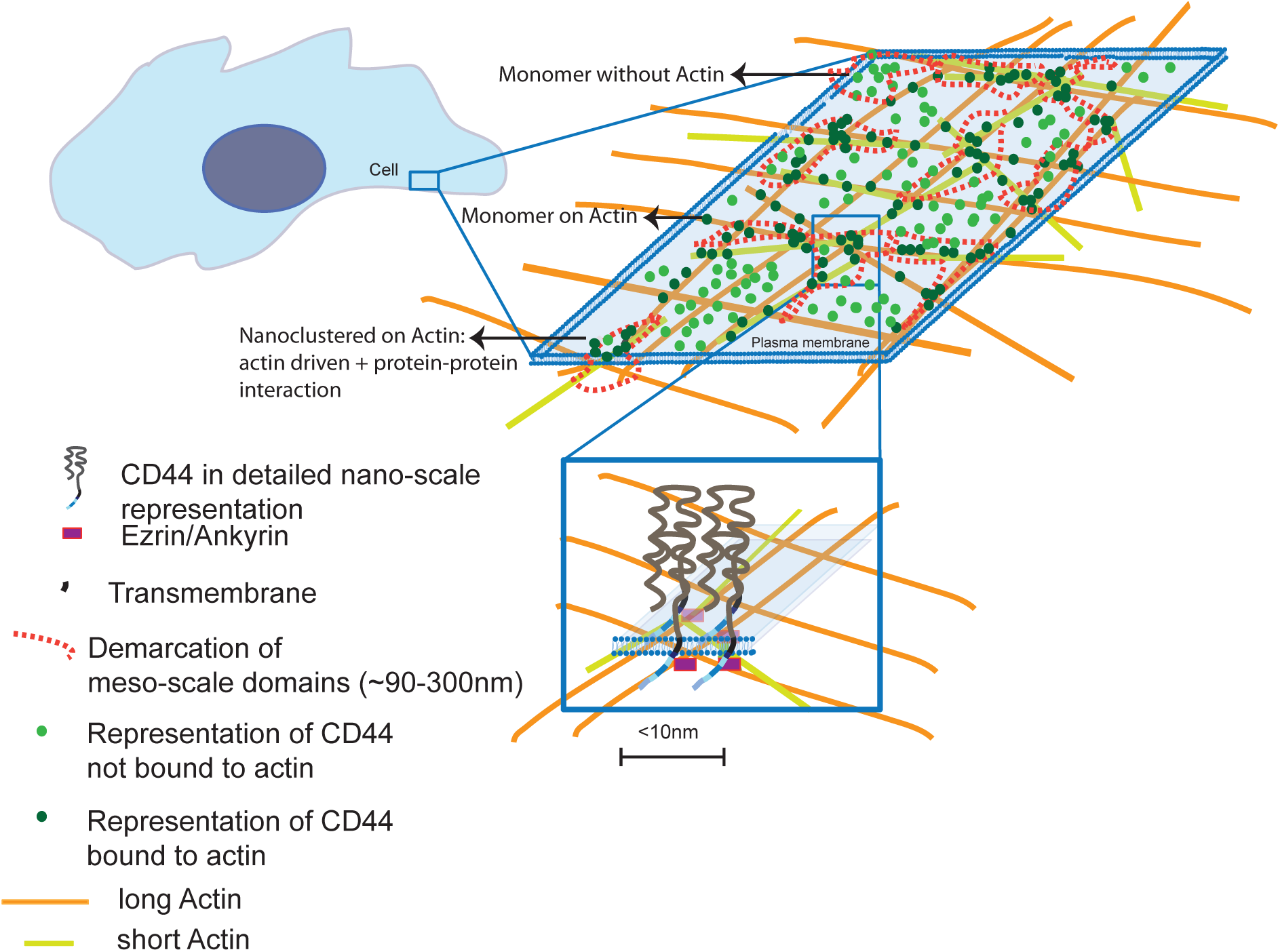
Proposed model for plasma membrane organization of CD44. In the cell membrane an ROI is outlined to show the distribution of monomers as well as clusters of CD44 receptors. Nanoclustered receptors are shown coupled to actin cytoskeletal elements by adaptors such as ezrin/ ankyrin (see zoomed-in nanocluster), interspersed with unattached CD44 molecules. The clusters of receptors are depicted as being driven by the action of formin polymerized actin filaments and myosin driven actin motility (molecules not depicted in the schematic). The meso-scale domains are CD44 localization hotspots identified in our experiment which are characterized by their close association with nanoclusters of the protein. The emerging meso-scale meshwork of the cell membrane receptor (depicted by the orange dotted line) may reflect the cytoskeletal meshwork juxtaposed to the plasma membrane.

The correlation between nano-scale and meso-scale organization of the protein and DC-SPT, reconciles the apparent heterogeneity in diffusion modes of molecules to confinement driven by clustering at spatially separated domains on the plasma membrane. From our meso-sale organization and SPT studies the regions on the membrane where the receptors are transiently confined/ temporarily arrested, correspond to regions of receptor co-localization as well as potentially, „localization hotspots’. These regions have an area ∼100-300 nm, outlining a fragmented meshwork-like pattern. Moreover, the timescale of turn-over of localization hotspots (Figure 7d) corresponds to the time scale of transient confinement of single molecules of CD44 (∼few (<3)sec; Figure 2). The receptor transiently associates with such regions and eventually unbinds to diffuse again, often guided by the underlying actin cytoskeleton-laid fences, until it encounters another suitable site at the membrane-cytoskeleton interface to be arrested again. Thus, we propose that our localization hotspots could correspond to the picket fences described earlier (Fujiwara et al., 2016; Murase et al., 2004).

To ascertain whether actin dynamics-driven mechanisms could template the nano and the meso-scale organization of CD44, we investigated the role of formin-nucleation based actin polymerization. As nanoclustering of CD44 is lost upon formin perturbation, we also observe concomitant lowering of the CD44 localizations detected on the underlying meshwork. This is reminiscent of the transmembrane domain of CD44 (CD44Tm-GFP) that cannot bind to actin. Additionally the meso-scale domain turnover is remarkably slowed down. This is consistent with previous studies that implicate the role of formin activity in the turnover of the underlying cortical actin meshwork (Fritzsche *et al*., 2013). These findings lead us to an important conclusion that meso-scale meshwork of CD44 arises as a consequence of the association of CD44 with the underlying actin cortex, and it is likely that the formin-mediated actin nucleation and turnover of the cortical actin meshwork contributes to the pool of dynamic actin necessary to template the nano-clustering of the protein as proposed previously (Chaudhuri *et al*., 2011). This also provides a natural explanation for the enrichment of CD44 nano-clusters along the meso-scale mesh which appears to mirror the cortical actin cytoskeleton mesh. At this time it should be noted that further experiments are necessary to prove the relationship between the cortical actin meshwork and the mesoscale meshwork of CD44.

Nano-clustering of CD44 is also abrogated upon removal of the cytoplasmic domain of CD44 (in Fig. 3f). This finding is further supported by cytoskeletal sensitivity of nanoclustering of the protein. The sensitivity of CD44 nanoclustering particularly to formin and ezrin perturbation is well aligned with the changes in CD44 diffusion upon similar perturbations, observed in SPT recently (Freeman *et al*., 2018). In that study, formin and ezrin mediated picketing function of CD44 had been implicated in regulating FcγRIIA dynamics and function in phagocytosis. Involvement of similar molecular machinery in nanoclustering as reported here strongly suggest that the picketed CD44 receptors are nanoclustered by the underlying dynamic actin filaments generated as a consequence of formin-driven actin polymerization, and driven by myosin activity.

In this study we have attempted to gain insights into specific interactions mediated by the ECD and ICD of CD44 in determining its diffusion and organization at the cell membrane. We find a strong correlation between nano-clustering potential and the tethering strength for the different truncation mutants of CD44 at the cell surface. Although removal of the ECD has little effect on the confinement radius of CD44, removal of the ICD from the mutant already lacking the ECD (CD44Tm-GFP) has a stronger effect on its confinement as well as localization on the meshwork at the mesoscale (Fig. 4d, Fig. 5b and Fig 5d). The ICD thus emerges as a stronger determinant for tighter confinement of CD44 at the membrane and as the domain that augments the registry of the mesoscale distribution with a meshwork pattern. Together with the result suggesting that the ECD deleted mutant still exhibits acto-myosin sensitive nano-clustering (Fig. S6), we believe that the meso-scale organization is templated on an underlying cortical actin mesh and serve to orchestrate the emergence of transient nanoclusters in its proximity.

The meshwork pattern that we observe may have a larger significance, since SNAP-CD44Tm-GFP and SNAP-CD59-GPI, proteins that are not directly coupled to actin, also exhibit meshwork-like appearance at the meso-scale. Spatially restricted diffusion patterns of the FcγRIIA which cannot interact with actin but associates with a CD44 defined mesh (Freeman *et al*., 2018), also exhibit non-random distribution at the meso-scale. This is likely to be mediated via lateral association of their membrane anchoring domains with actin-binding membrane pickets, or confinement within membrane compartments demarcated by picketing proteins. These data supports the picture of a tightly coupled actin-membrane composite where even proteins that do not couple to actin are impacted by the patterning of the underlying meshwork.

With further sophistication of imaging and analysis methods, the correlation of cartography and anisotropy can be studied with higher temporal resolution. While our study is currently restricted to cytoskeletal interactions of CD44, there remains scope for detailed analysis of the influence of the exo-plasmic interactions with molecules such as galectins and HA. Simultaneous imaging of signaling and cytoskeletal adaptors along with CD44 can open up possibilities for exploring potential outside-in (ligand binding can lead to signaling adaptor recruitment) as well as inside-out signaling (ankyrin binding can influence hyaluronic acid binding (Zhu & Bourguignon, 2000)) at the nano and meso-scale domains. Since CD44 is implicated in processes such as metastasis, phagocytosis or lymphocyte rolling (Donatello et al., 2012; Hanke-roos et al., 2017; Hill et al., 2006; Vachon et al., 2018), they provide physiologically relevant scenarios where local and global organization of CD44 may have an impact on relevant physiological scenarios.

We believe that the spatial organization of CD44 determined by the dynamic remodeling of the actin cytoskeleton, define dynamic fences that partition the receptor in different regions of the cell membrane. These fences have been implicated in the phagocytic function of FcγRIIA, and the endocytosis of DC-Sign receptor, which are receptors that do not exhibit direct interaction with the actin cytoskeleton (Freeman et al. 2018; Torreno-pina et al. 2014). In conclusion, our approach and findings provide a multi-scale view of organization of a trans-membrane protein at the cell membrane, revealing a hierarchical framework where actomyosin-driven nano-clusters emerge in close association with an underlying dynamically remodeling meso-scale meshwork, enabling the cells to spatio-temporally regulate receptor organization.

## Materials and Methods

### Plasmids, cell lines and antibodies

CD44-GFP, CD44ECDTm-GFP, CD44TmICD-GFP cloned in p-EGFP N1 vector were gifts from Rob Parton in University of Queensland Australia. CD44ΔERM-GFP, CD44ΔAnk-GFP, CD44ΔEA-GFP and CD44Δ15-GFP constructs were generated by site directed mutagenesis using CD44-GFP as the template in the same backbone. SNAP and FR tagged CD44 constructs were designed and cloned into a lentiviral pHR transfer backbone and cloned between MluI and BamHI/ NotI sites using Gibson Assembly method. All constructs were sequenced and verified using appropriate primers (Table 2). SNAP CD59 GPI was obtained from Addgene (Addgene #50374). Sequences and primer sequences will be made available upon request. Cell line expressing FR-CD44TmICD and FR-CD44Tm were generated by transfecting and selecting transfected cells by staining for FR expressing cells with anti folate receptor MOV19 antibody using fluorescence assisted cell sorting (FACS). CHO (Chinese Hamster Ovary) cells were cultured in Ham’s F12 media (HiMedia, Mumbai, India); MCF-7, COS-7 (African green monkey kidney cells) and MEFs (Mouse embryonic fibroblasts) were cultured in DMEM High-Glucose (Gibco^TM^, 21720-024). The media was supplemented with fetal bovine serum (FBS) (Gibco^TM^, 16000044) and a cocktail of Penicillin, Streptomycin, L-Glutamine (PSG) (Sigma, G1146-100ml). MEFS, MCF-7, COS-7 or CHO cells were seeded sparsely and grown for 2 days on 35mm cell culture dishes fitted with a glass bottom coverslip for imaging. Cells were transfected with the different CD44 plasmids, 12-16 hrs before imaging, using Fugene® 6 Transfection reagent (E2692, Promega).

### Antibody labeling and expression level estimation

Endogenous and over-expressed CD44 on the cell surface in the different cell lines, plated on cover-slip bottom 35mm dishes, after 2 days of plating, were labeled using IM7 antibody (14-0441-82, eBioscience™) on ice for 1hour followed by incubation with anti-Rat secondary antibody tagged to Alexa 633 (A21094, Life technologies) on ice for 1hour. The antibodies were diluted in 10% FBS containing culture media (DMEM). The cells were washed and imaged in HEPES buffer and imaged using a 20X objective on a spinning disc microscope. Mean intensity from ROIs drawn around cells was quantified using ImageJ.

### Actomyosin perturbation

Blebs were generated using 14μM Jasplakinolide (Thermo Fischer, Invitrogen, catalog no.: J7473) for 15min. Formin perturbation was carried out using 10-25μM SMI-FH2 (Calbiochem, catalog no.: S4826-5MG) for 15min-1 hour based on experimental requirement. Arp2/3 inhibition was carried out using 200uM CK-666 (Sigma Aldrich, Catalog no: SML0006 5MG) treatment for 3 hours. Ezrin perturbation was carried out using the inhibitor NSC668394 purchased from EMD Millipore, (Cat. No: 341216-10MG). Cells were treated with 25μM of the drug, for 1 hour. Myosin II perturbation was carried out using a cocktail of ML-7 (Sigma Aldrich, Catalog no.: I2764) and Y27632 (Sigma Aldrich, Catalog no: Y0503-1MG) or H1152, purchased from Tocris (Ctalog no.: 2414). Cells were treated with a cocktail of the ML-7 and Y27632/ H1152 at a final concentration of 20 μM of each, for 1hour. Due to the reversible nature of the drugs acting on the target, imaging was carried out in the presence of the drug except in the case of Jasplakinolide treatment. All drug treatments were carried out in HEPES buffer saline containing 2mg/ml glucose at 37 degrees for the indicated time periods.

### STORM sample preparation and imaging

CHO cells were plated on a 8-well Lab-Tek #1 chamber slide system (Nunc) at a density of 30000 cells/well. Cells were incubated at 37°C for 24 hours. After incubation, the samples were fixed with 4% paraformaldehyde in PBS at room temperature for 20 min. After fixation, blocking solution (3% w/v BSA in PBS) was applied for 30 min. Cells were labeled with rat-anti-mouse-anti-CD44 primary antibody (Clone KM114, BD Pharmingen #558739) at a concentration of 5ug/ml for 1 hour at room temperature. The corresponding secondary antibody (anti-rat) was tagged with Alexa Fluor 647 (Invitrogen) as a reporter and with Alexa Fluor 405 as an activator. The secondary antibody was incubated for 1 hour at room temperature. Cells were stored in 1% PFA in PBS. The STORM buffer used was the same of Gómez-Garcia et al (Gómez-García *et al*., 2018): Glox solution (40 mg/ml Catalase [Sigma], 0.5 mg/ml glucose oxidase, 10% Glucose in PBS) and MEA 10mM (Cysteamine MEA [Sigma Aldrich, #30070-50G] in 360mM Tris-HCl). The imaging for STORM on endogenous CD44 from top surface in CHO cells is from one experiment.

In order to study the nearest-neighbour distribution of clusters, we identified the clusters of localizations based on intensity (i.e. high density of localizations) and determined the position of the center of mass. With this information we calculate the NND for the experimental set. For the simulations, we take the same identified clusters (keeping their size) and reshuffle them in space. We repeat this process many times (100 times) to get more robust information on the simulated NND.

### Live cell imaging for fluorescence emission anisotropy and cartography experiments

All live imaging were interchangeably carried out, based on requirement, in one of the following set-ups: (1) confocal Spinning disk microscope (for imaging blebs in 3-D) equipped with a Yokogawa CSU-22 unit and 100x, 1.4NA Nikon oil objective, Andor technologies laser combiner emitting 488nm and 561nm wavelength amongst others and Andor ixon+897 EMCCD cameras. Images were acquired using Andor iQ2 software (2) TIRF microscope set-up equipped with Nikon Eclipse Ti body, a 100X, 1.45NA Nikon oil objective, photometrics Evolve EMCCD cameras, an Agilent laser combiner MCL400 (Agilent technologies) whose 488, 561nm and 640nm excitation wavelengths were used as necessary and µManager for image acquisition (3) TIRF microscope setup equipped with Nikon TE2000 body, a 100X, 1.49NA Nikon oil objective, EMCCD Cascade 512 cameras (photometrics Inc., Tuscon, USA), home-built laser combiner equipped with 488 and 561nm lasers, and Metamorph^TM^ /µManager for image acquisition). Wherever necessary, live imaging was performed in a temperature controlled stage-top incubator chamber with immersion thermostat, ECO Silver, from Lauda Brinkmann.

### Fluorescence Emission Anisotropy measurements

We measure emission anisotropy of our protein of interest by labeling them with GFP or PLB, both of which are suitable for fluroscence anisotropy measurement to report on Homo-FRET(Sinnecker *et al*., 2005; Ghosh *et al*., 2012). Cells were imaged in HEPES buffer containing 2mg/ml glucose on an inverted TIRF microscope using polarized excitation light source. Emission was split into orthogonal polarization components using a polarization beam splitter and collected simultaneously by two EM CCD cameras to detect polarization of emitted fluorescence. Fluorescence emission anisotropy measurements were interchangeably carried-out, based on requirement, in one of the dual camera equipped imaging systems described before. Steady state fluorescence emission anisotropy was calculated as elaborated in Ghosh et al, 2012.The absolute value of anisotropy is a function of the effective numerical aperture of the imaging system (Ghosh *et al*., 2012). Since the effective numerical aperture is determined by combinatorial effect of individual lenses in the light path of the microscope system, the absolute anisotropy value of the same protein varied from one system to another. Also, since the different experiments reported here have been conducted over several years, absolute values of anisotropy for the same constructs would have varied based on the status of the optics in a given microscope system. Hence, the measurements typically contained an internal control for sensitivity of anisotropy change, which was generally measurement of the extent of anisotropy change between the wild type CD44-GFP and CD44ECDTm-GFP (or CD44-TmICD-GFP and CD44-Tm-GFP).

### Fluorescence anisotropy image analysis

Fluorescence emission anisotropy of GFP and PLB tagged proteins was calculated using images from the two cameras which were individually background corrected and the perpendicular image G-Factor corrected (Ghosh *et al*., 2012) to rectify effects of inherent polarization bias of the imaging system using imaging software: ImageJ or Metamorph^TM^. 20X20 or 30X30 pixel ROIs were drawn to sample the cell membrane and anisotropy values from the specified regions were obtained. Anisotropy maps were generated after aligning the images from the two cameras and calculating pixel-wise anisotropy value as described in (Ghosh *et al*., 2012), using a custom code written in MATLAB (The Mathworks, Natick, MA). Code will be available upon request. For data plotting, intensity was binned for appropriate intensity range and each data point represents mean and error bar represents standard deviation of anisotropy corresponding to the intensity bin. We ensured that data comparisons were done between conditions across similar intensity range. Intensity range chosen was decided based on different microscope properties, especially the bit depth and noise levels of the cameras. For representation calculated anisotropy values from the intensity images of the parallel and perpendicular cameras have been plotted on the Y axis as a function of the expression level, which is described as „Total intensity in arbitrary units’ on the X axis. Here the total intensity is computed as a summation of the intensity recorded in the parallel image and two times the intensity recoded in the perpendicular image as described in Ghosh et al., 2012.

### Labeling of SNAP tagged-CD44 membrane receptors

MEFS, COS-7 or CHO cells were seeded sparsely and grown for 2 days on 35mm cell culture dishes fitted with a glass coverslip at the bottom. Cells were transfected with the different SNAP tagged CD44 plasmids 16-18hrs prior to the experiment using Fugene® 6 Transfection reagent. Labeling were done with SNAP tag specific photo-stable fluorescent probes SNAP alexa 546, SNAP-surface® 549 (ex/ em: 560/575 nm, purchased from NEB, USA) or JF646-SNAP ligand ( ex/ em: 646/664 nm) by incubating for 10 min at 37° C using dilution of 30 nM (for single particle experiments) and 50-100nM (for cartography experiments) with 10% serum containing F12 medium and then washed extensively with glucose-M1 buffer (150 mM NaCl, 5 mM KCl, 1 mM CaCl_2_, 1 mM MgCl_2_, 20 mM HEPES, pH 7.3; supplemented with D-glucose at 2 mg/ml) to get rid of free dyes. The dyes were chosen to ensure they are spectrally different from GFP with minimum bleed-through. Dual color labeling was done with JF549-cpSNAP ligand ( ex/ em: 549/571 nm) and JF646-SNAP ligand ( ex/ em: 646/664 nm) fluorophores by incubating for 10 min at 37° C with F12 serum medium at a mixed concentrations of 50 nM and 150 nM for the respective dyes. Singly or dually labeled cells were subsequently washed and imaged at 37° C in presence of HEPES buffer containing 2mg/ml glucose.

### Single particle tracking

Video imaging of single fluorescent receptors on cell membrane were performed using a home built, total internal reflection fluorescence (TIRF) microscope equipped with a Nikon Eclipse Ti body and a Nikon 100X Apochromat 1.49 NA objective, with a C-MOS sensor based high-speed camera (FASTCAM-SA1, Photron, Tokyo, Japan; (Shibata *et al*., 2012; Hiramoto-Yamaki *et al*., 2014; Komura *et al*., 2016)) coupled to a two-stage micro-channel plate intensifier (C8600-03, Hamamatsu Photonics, Hamamatsu, Japan) by way of an optical-fiber bundle. Single molecules were observed at 16.7 ms (60 fps) temporal resolution with excitation laser of 561 nm of power density ∼ 2.43 kW/cm^2^, with a FWHM (full-width-half-maximum) of 333 ± 13 nm and pixel size of 54 nm, in presence of an additional 1.5x lens in front of the camera. The localization precision was estimated to be ± 28 nm. The precision was measured after immobilizing CD44 labeled with SNAP-surface® 549 fluorescent probe, on MEFS cells, by fixing the cell membranes with 4% paraformaldehyde and 0.1 % glutaraldehyde for 60 min at room temperature (Tanaka *et al*., 2010). The precision was determined by fitting the centroid position from single molecules using a 2D Gaussian function and calculated from radial standard deviation δ_r_ = (δ_x_ * δ_y_)^1/2^ ≈ δ_x_ ≈ δ_y_ of x, y-coordinates over time. Tracking of membrane molecules (x- and y-coordinates) were determined using C++ based computer program as described previously (Fujiwara et al., 2002; Koyama-honda et al., 2005). The mean-squared displacement (MSD) for every time frame for each trajectory was calculated as per following equation:

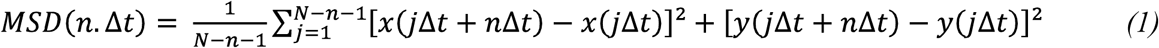

Where, Δt is the time increment, N is the number of frames of the trajectory, n is the number of time increments, and x and y represent the particle coordinates. Then, the microscopic diffusion coefficients (D_2-5_) of individual trajectories were calculated through a linear fit performed at short time lags (n = 2-5) using the equation

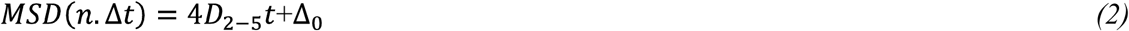

Where, the MSD intercept at zero time lag, Δ_0_ is associated to the localization precision.

### Mobile fractions and temporal confinement detection

Temporal confinement or Temporary Arrest of Lateral diffusion (TALL) were analyzed defining parameters of detection circular radius and threshold residence time by using the algorithm developed by *Sahl et al*. (Sahl et al., 2010). Theoretically, simulated randomly diffusive trajectories show false TALL of ∼5% of total trajectory lengths. Therefore, the detection of circular radius was set, based on calculating average diffusion coefficient (0.3 µm^2^/sec) of mobile fractions of CD44 and probability of temporal confinement < 5% during Brownian motion within, 10 frames of 16.7 ms exposure. In escape probability method, the probability P(r,t) that a particle diffusing with the diffusion coefficient D, remains confined within the circle of radius r and the time interval t can be expressed as:

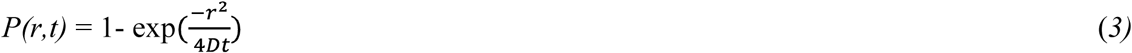

Data represented is pooled from two different replicates, which individually exhibited similar trend. Choice of cells from which data has been represented was made based on optimal labeling density and flatness of membrane morphology since imaging has been done in the TIRF mode.

Analysis was extended to determine the discrete probability density *P(Δr^2^,Δt)* by cumulative square displacements, which will represent a sequence of spatial positions *r(t)* separated by variable time lags *Δt.* The cumulative probability *P(Δr^2^,Δt)* is defined by equation (3), where α is the time fraction of characteristic free diffusion with coefficient D, 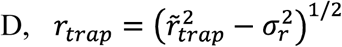 corresponds to the trapping radius of the particle and _r_ is the experimental localization error (Sahl *et al*., 2010).

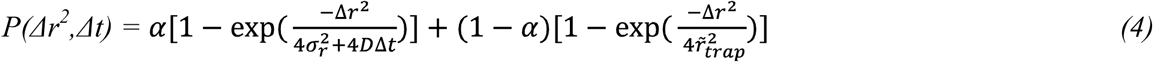

Probability density *P(Δr^2^,Δt)* observing for long steps were corrected with overlap integral of two circles with radius R by *P_track_ (Δr,R)* as described in *(Sahl et al. 2010)*. Here we computed *P(Δr^2^,Δt)* with increments *Δ(Δt)=* 16.7 ms and *Δ(Δr^2^)=* 50 nm^2^.

### Dual color trajectory analysis

Inter-molecular separation distance between CD44 molecules labeled with JF549 (green) and JF646 (red) dyes was determined from the centroid locations of their dual color pair trajectories within boundaries ranging from 25nm to 500nm radius using C++ based computer program WinCol (Koyama-Honda *et al*., 2005). Measurements were done with excitation lasers of 561 nm and 642 nm of power density ∼ 2.43 kW/cm^2^ and ∼ 4.06 kW/cm^2^ respectively, and detecting signals simultaneously by two cameras after splitting emission signals using a 561/647 dichroic mirror (Chroma Technology, 625DCXR) with corresponding emission band pass filters 593 ± 43 nm and 685 ± 40 nm. Localization accuracy of JF549 and JF646 dyes are ± 29 nm and ± 33 nm, respectively while pixel size at image plane is 54 nm. Videos of the flipped green channel were used to generate randomly encountered co-localizations. Co-localization was defined when intermolecular distances were 200 nm for a minimum of 3 consecutive frames. The displacement between co-localized frames was then calculated. The displacement and step size distribution were thereafter compared with transiently confined frames, trajectories of mobile fractions and all frames. Photo-bleaching analysis from individual spots of fluorophores did not reveal any significant bleaching in the timescale reported for the lifetime of co-localization of the protein (Fig. S2d). Data represented is pooled from two different replicates, which individually exhibited similar trend. Choice of cells from which data has been represented was made based on optimal labeling density and flatness of membrane morphology since imaging has been done in TIRF mode. Total number of trajectories analyzed: SNAP-CD44-GFP = 27856, SNAP-CD44Tm-GFP= 7516.

### Generation of cartography

Cartography maps were generated from movies (1000 frames, 10 fps) recorded in TIRF mode as explained in the previous section, using sub-saturation labeling conditions (50nM-100nM). Identification of single molecules essentially corresponds to the identification of individual fluorescent spots at each given time frame. For this, we apply two criteria: First, the spots should have a size that is limited by diffraction, i.e. this corresponds to the PFS of the microscope. Second: the intensity of each spot should be higher than the surrounding background. The localization precision of each individual spot is given by the number of counts on that spot, which in the case of our videos corresponds to ∼ 20nm. The spatial (x,y) coordinates of the labeled membrane receptors (for each of the constructs investigated) were thus retrieved from each frame using a MATLAB routine based on Crocker (1996)(Crocker and Grier, 1996), with sub-pixel accuracy. Finally, all the receptor coordinates of all frames were collapsed into a single image, the so-called the cartography map. With this approach, not only one can access the nanoscale organization of the labeled receptor, but also the mesoscale organization without the need of reconnecting trajectories (Torreno-pina et al., 2014). Cartography maps were also generated in different time windows, typically by integrating the localizations over 40 (Fig. 2) or 20 frames (Fig. 1e, Fig. 5). Experiment to obtain cartography maps of the receptor and the mutants have been conducted at least twice in MEFs and once in COS-7 cells. Formin perturbation and mesoscale organization imaging has been done at least twice and the represented experiment here is done in COS-7 cells. SNAP-CD44-GFP cartography in CHO cells and GPI mesoscale organization experiment has been conducted once.

### Analysis of the cartography maps

Since the cartography maps are generated from localizations obtained as a function of time, their evolution is dynamic. Therefore, we restricted our analysis to time windows of 2s by collapsing all the localizations from sequential 20 frames, into a single, less crowded, cartography image. Confinement areas were identified using the MATLAB routine DBSCAN (Density-Based Spatial Clustering of Applications with Noise) with settings (epsilon = 1.0 and MinPts = 10). Finally, we defined the confinement area as the area occupied by a cluster of localizations.

For the time-evolution analysis of the meso-scale domains, the time windows correspond to 2 seconds, i.e. 20 frames. Initially clusters are defined at the time window 0, (frames within f0 and f0+19). Then, since we slide the window through the cartography map, at each time window we move 100 ms in the cartography.

### Analysis of the interleaved anisotropy and cartography maps

In order to compare the cartography maps with the anisotropy images, we performed interleaved anisotropy imaging together with high density SPT, generating one anisotropy image before starting SPT, a second anisotropy at frame 500 of the SPT recording and a final one once the SPT recording was finished (after frame 1000). To reduce temporal variations on both the anisotropy and cartography maps, we focused on anisotropy images at the corresponding frame 500 of the SPT movie. The anisotropy image was divided into small ROIs (22-by-22 pixels, with a pixel size of 106nm). This was done in order to select only those regions were the plasma membrane is completely flat and therefore the anisotropy arises exclusively from the lateral distribution of the labeled receptors. In addition to this, for each ROI, we classified each pixel of the anisotropy map into 3 three groups: low anisotropy (Low A), median anisotropy (Medium A) and high anisotropy (High A).

We then took the localizations between frames 480 and 520 of the SPT movie and generated a cartography map for each of the ROIs. We identified the clusters of localizations using the MATLAB routine DBSCAN with settings (epsilon = 1.0 and MinPts = 10). With the localizations belonging to clusters, we assigned to each of them an anisotropy value corresponding to their location in the anisotropy ROI and classified them within the three groups. Simultaneously, we randomly distributed the same number of localizations on the anisotropy ROIs and also assigned their corresponding anisotropy value and posterior classification. Comparative anisotropy-cartography analysis has been done from an experiment with COS-7 cells where localization and GFP based FRET information was obtained using dual cameras at specific intervals during acquisition of single molecule localization time series of the SNAP tag fluorophore.

### Statistical Analysis

Differences between anisotropy distributions between control and treatment were tested using non-parametric Mann-Whitney test or KS test. Number of fields/ cells imaged is mentioned against each experiment. The anisotropy and cartography data shown here is from one representative experiment. Each experiment has been conducted at least twice unless otherwise mentioned. Quantification from cartography and SPT experiments has been tested for significance using Kruskal Wallis test along with post hoc Tukey-Kramer test and Wilcoxon sum rank test of Matlab unless otherwise mentioned.

## Acknowledgements

We would like to thank Rob Parton for sharing with us the CD44-GFP, CD44ECDTm-GFP and CD44TmICD-GFP constructs, which were originally used in Mrass et al., 2012. The JF dyes used in SPT studies were a generous gift from Luke Lavis, HHMI Janelia Research Campus. We would like to thank Suvrajit Saha for contributions to the initiation of the project. We thank Central Imaging and Flow facility of NCBS for enabling us to use their equipment and Divya Gowda and H. Krishnamurthy for their help. We thank Marcus J. Taylor for helping in reagent development pertaining to the SNAP tagged constructs. We would like to thank Chaitra Prabhakara and Sanjeev Sharma for help with data representation, revision and editing the manuscript. M.F.G.-P acknowledges funding from the Fundació Privada Cellex, Generalitat de Catalunya through the CERCA program, Spanish Ministry of Economy and Competitiveness (“ Severo Ochoa” Programme for Centres of Excellence in R&D (SEV – 2015 –0522) and FIS2017-89560-R) and from European Union H2020-ERC grant 788546-NANO-MEMEC. N.M. acknowledges funding from the European Union H2020 under the Marie Sklodowska-Curie grant 754558-PREBIST. C.M. acknowledges funding from the Spanish Ministry of Economy and Competitiveness and the European Social Fund through the “Ramón y Cajal” program 2015 (grant no.RYC-2015-17896) and the “Programa Estatal de I+D+i Orientada a los Retos de la Sociedad” (grant no. BFU2017-85693-R), and from the Generalitat de Catalunya (AGAUR Grant No. 2017SGR940). A. K. acknowledges support in part by Grants-in-Aid for scientific research Kiban S (16H06386) from the Japan Society for the Promotion of Science. TK.F. acknowledges „Grants-in-Aid for Scientific Research from the Japan Society for the Promotion of Science (Kiban B (16H04775))’. KS acknowledges ‘Grants-in-Aid for Scientific Research from the Japan Society for the Promotion of Science (Kiban B (18H02401))’.S.M. acknowledges JC Bose Fellowship from DST (Government of India), a collaborative grant from HFSP (RGP0027/2012 with M.F.G.-P) and Wellcome Trust-DBT Alliance Margadarshi fellowship (IA/M/15/1/502018).

