## Supplementary figures and legends for "Dynamic actin-mediated nano-scale clustering of CD44 regulates its meso-scale organization at the plasma membrane"

**Supplementary figure legends**

**Figure S1: Non-random distribution of CD44.** (a) Cartography map of SNAP-CD44-GFP expressed in COS-7 cells, obtained at sub-saturation labeling conditions (~50-100 nM); (x,y) coordinates obtained from 1000 frames collapsed in a single map with a zoomed-in ROI . (a') Cartography construction of (x,y) coordinates in the marked ROI in (a) from 50 consecutive frames obtained at two different experimental time windows, between 30s-35s (magenta, left) and 90s-95s (green, centre) and merged image indicating persistent domains in white (right). (b) Plot depicting distribution of confinement area in MEFs (Sub-plotted for representation, from Fig.5). (c) An example cartography snapshot of SNAP-CD44-GFP expressed in CHO cells. (d) Representative STORM images of endogenous CD44 acquired from the top membrane in CHO cells indicating the mesh-like pattern with white dotted lines. (e) Nearest neighbor distance(NND) plot of CD44 clusters detected in STORM compared to randomised simulation depicting the heterogeneous nature of clustered CD44 distribution at the membrane (by Wilcoxon rank sum test, $p<e^{-18}$). (f) Cartography of SNAP-CD59 GPI obtained under similar labelling conditions as CD44 depicted at two different magnifications.

**Figure S2: Colocalization of CD44 by DC-SPT.** (a) A typical trajectory of SNAP-CD44-GFP expressed in MEFs, depicting regions of free diffusion in cyan as well as regions of confinement in green along with zoomed-in view of a region of transient confinement. (b) Distribution of lifetime of transient confinement for SNAP-CD44-GFP. (c) Occurrence of co-localization events (i.e., inter-particle distances < 200nm) per unit area, as a function of inter-particle distance, compared to randomized trajectories (n=7cells, p<0.0011). (d) Intensity traces of individual spots of JF-646 SNAP ligand (top in red) and JF549 cpSNAP ligand (bottom in green) exhibit minimal photo-bleaching in the time-scale of co-localization between the two differently labeled fluorophores as reported in Fig. 1f (e and e') 3D-trajectory of the full length SNAP-CD44-GFP and the SNAP-CD44Tm-GFP depicting motion of the dual-color (coded in green, JF549-cpSNAP ligand and red, JF646-SNAP ligand) labeled receptors, sampled, over a period of 650 milliseconds. (f) Plot depicts traces of inter-particle distance measured for full length CD44 and the trans-membrane domain alone over entire trajectories. Dotted line indicates inter-particle distance of 200nm. (f') Comparison of inter-particle distance distribution of the full length CD44 (SNAP-CD44-GFP) and the CD44 trans-membrane domain (SNAP-CD44Tm-GFP). Here (f and f') the data has been represented for trajectories from a single cell. Pooled data points from all cells are not represented here to avoid over-crowding.

**Figure S3: Anisotropy measurements upon photo-bleaching of FR-CD44 chimera in CHO cells and effect of intra-cellular domain on CD44-GFP anisotropy in different cell types.** (a) Schematic showing folate receptor (FR) tagged CD44 constructs that can be exogenously labeled with a fluorescent folate analogue ($PLB^{TMR}$: $N^{\alpha}$-pteroyl-$N^{\varepsilon}$-$Bodipy^{TMR}$-L-lysine) to enable anisotropy imaging upon photo-bleaching. (b) Montage shows snap-shots of intensity and anisotropy images of FR-CD44TmICD and FR-CD44Tm expressed in CHO cells, during different times after initiation of photo-bleaching. (c) Quantification of anisotropy as a function of the extent of photo-bleaching reveals a distinctly higher slope for FR-CD44TmICD indicating that it is clustered to a greater extent than the FR-CD44Tm. Note the starting anisotropy of the FR-CD44TmICD is also much lower that FR-CD44-Tm, consistent with the increased extent of clustering of the FR-CD44-TmICD. (n= 2 fields for each in the representation) (d-f') Graphs show comparison of CD44-GFP and CD44ECDTm-GFP anisotropy versus intensity distributions in the (d,d') COS-7 cells (CD44-GFP(n) = 18 fields, CD44-ECDTm-GFP (n) = 14 fields) and (e, e') MCF-7 cells, both of which have low levels of membrane CD44 (CD44-GFP(n) = 16 fields, CD44-ECDTm-GFP (n) = 18 fields) and (f, f') CD44 expressing and HA secreting MEFs(CD44-GFP(n) = 10 fields, CD44-ECDTm-GFP (n) = 13 fields) respectively. In each of the cell lines the anisotropy of the CD44-GFP is significantly lower than the comparative values obtained from CD44ECDTm-GFP ($p< 10^{-55}$), indicating significantly higher clustering potential of the full length construct compared to cytosolic tail deleted protein, regardless of the cell line in which it is expressed. (g) plot depicting relative levels of cell surface CD44 stained by anti-CD44 antibody in the different cell lines used in the study and comparison of the level of over-expression achieved in CD44 deficient cells with MEFs that endogenously express the protein (MEFs un-transfected (n)= 87 cells, MCF-7 un-transfected (n) = 102 cells, MCF-7 CD44GFP (n)= 32 cells, COS7 un-transfected (n)= 70 cells, COS7 SNAP-CD44-GFP (n) = 51cells, COS7 CD44-GFP(n)= 102 cells. $p<10^{-3}$).

**Figure S4: Comparative analysis of dynamics of different truncated mutants of CD44 by SPT.** (a) More sample trajectories of the SNAP tagged CD44 constructs. (b) Quantification of discrete probability density P ($\Delta r^2$, $\Delta t$) by cumulative square displacements for the indicated SNAP tagged CD44 constructs. (c) Mobile fraction analysis of the three proteins in MEFs calculated by using discrete probability density P ($\Delta r^2$, $\Delta t$) by cumulative square displacements. (n=6) Difference between distributions was tested for significance using Kruskal-Wallis and post hoc test with Tukey-Kramer. Comparison of (d) mobile fractions and (e) confinement radius (r_trap) between SNAP tagged CD44 and its mutants, in COS-7 cells. SNAP-CD44-GFP and SNAP-CD44TmICD-GFP (n) =5 and SNAP-CD44Tm-GFP (n) = 4 cells, with differences between distributions tested by Mann-Whitney test. Number of trajectories: SNAP-CD44-GFP = 1155, SNAP-CD44TmICD GFP= 1026, SNAP-CD44Tm-GFP = 802.

**Figure S5: Analysis of the meso-scale organization of the different constructs in CD44-deficient COS-7 cells.** (a) Cartography maps of the indicated SNAP tagged CD44 constructs expressed in CD44-null COS-7 cells, obtained by imaging sub-saturation labeled (50-100 nM) at 10fps and accumulating the spatial coordinates of individual molecules over 2s period. (b) Quantification of the confinement areas for the different constructs over 2s. Black lines correspond to the mean value. (c) Normalized fraction of confinement areas/area for the three different constructs. A significantly lower number of confined regions per unit area are observed in the SNAP-CD44Tm-GFP as compared to the full length receptor or the mutant lacking the extra-cellular domain. This can also be directly inferred from the cartography maps where the localizations are more tightly bound for the full length receptor and much more disperse in the case of the SNAP-CD44Tm-GFP mutant. (d) Fraction of localization events belonging to the meshwork for the wild type and the mutant construct in COS-7 cells. SNAP-CD44-GFP (n) = 9 cells; SNAP-CD44TmICD-GFP (n) = 6 cells; SNAP-CD44Tm-GFP(n)= 5 cells. Difference between distributions was tested for significance using Kruskal-Wallis and post hoc test with Tukey-Kramer. (b): SNAP-CD44-GFP & SNAP-CD44TmICD-GFP: $p<e^{-9}$→ ns; SNAP-CD44-GFP & SNAP-CD44Tm-GFP: $p <e^{-8}$; SNAP-CD44TmICD-GFP & SNAP-CD44Tm-GFP: $p<e^{-9}$. (c): SNAP-CD44-GFP & SNAP-CD44TmICD-GFP: $p<0.05$; SNAP-CD44-GFP & SNAP-CD44Tm-GFP: $p < 0.05$; SNAP-CD44TmICD-GFP & SNAP-CD44Tm-GFP: $p=0.997$. (d): SNAP-CD44-GFP & SNAP-CD44TmICD-GFP: $p=0.5386$→ ns;

SNAP-CD44-GFP & SNAP-CD44Tm-GFP: $p = 0.0855$ → ns; SNAP-CD44TmICD-GFP & SNAP-CD44Tm-GFP: $p< 0.005$.

**Figure S6: Sensitivity of wild type and CD44 without ECD to actin dynamics.** (a) CD44TmICD-GFP is sensitive to Actin perturbations as observed in Jasplakinolide treated cells. CD44TmICD-GFP anisotropy on blebs is higher than flat membrane of untreated cells as depicted in the plot. (In DMSO treated, n=5 fields; Jas treated, n=19 fields; $p<10^{-7}$). (b) CD44TmICD-GFP is sensitive to perturbations of ezrin function, consistent with the effects of ezrin perturbation on the full length protein. (Con (n) = 15 fields, Treatment (n) = 21 fields; $p<10^{-50}$, by Mann Whitney test) (c) CD44TmICD-GFP nanoclustering is sensitive to actomyosin perturbations as depicted by the higher anisotropy of the cells treated with the cocktail of inhibitors (MLH) to prevent myosin II regulatory light chain phosphorylation, compared to untreated control cells ((n)= 11 fields for both control and treatment; $p<10^{-50}$). (d) CD44-GFP nano-clustering is mildly sensitive to Arp2/3 perturbation as depicted by the higher anisotropy value of CK666 treated cells (CK666 200µM, 2 hours; Con (n) = 12 fields, Treatment (n) = 12 fields, $p<10^{-5}$).

**Figure S7: Role of multiple cytoskeletal adaptor binding domains in the CD44 tail in nano-clustering across different cell types.** (a, a' and a'') Plots show intensity versus anisotropy distributions of the CD44 mutants in HA secreting MEFS (a; Distribution of anisotropy values were tested for significance using Mann-Whitney test and $p<10^{-75}$ was obtained for CD44-GFP and ECDTm-GFP; CD44-GFP (n) = 21 fields, CD44-ECDTm-GFP (n) = 15 fields, CD44-Δ15GFP (n) = 16 fields, CD44-ΔEA-GFP (n) = 14 fields, CD44-ΔERM-GFP (n) = 12 fields), HA deficient CHO cells (a'; CD44-GFP (n) = 16 fields, CD44-Δ15GFP (n) = 16 fields, CD44-ΔERM-GFP (n) = 16 fields, CD44-ΔEA-GFP (n) = 14 fields, CD44-ΔAnk-GFP (n) = 18 fields) and COS-7 cells which exhibit low surface levels of CD44 (a''; Distribution of anisotropy values were tested for significance using Mann-Whitney test and $p<10^{-184}$ was obtained for CD44-GFP and CD44ECDTm-GFP; CD44-GFP (n) = 17 fields, CD44-ECDTm-GFP (n) = 12 fields, CD44-Δ15GFP (n) = 21 fields, CD44-ΔEA-GFP (n) = 16 fields, CD44-ΔAnk-GFP (n) = 16 fields) The difference in anisotropy distribution between wild type and other mutants such as CD44ΔERM-GFP/ CD44ΔAnk-GFP/ CD44ΔEA-GFP/ CD44Δ15-GFP is minimal. The data is from one representative experiment.

**Supplementary Video1:** Video depicting motion of single particles of SNAP-CD44-GFP labeled with two different dyes (coded in green, JF549-cpSNAP ligand, and red, JF646-SNAP ligand) acquired for 400 frames at 16ms exposure time.

**Supplementary Video 2:** Video depicting a track of two differently labeled SNAP-CD44-GFP (coded in green, JF549-cpSNAP ligand, and red, JF646-SNAP ligand) showing co-localization of the protein over multiple frames.

**Supplementary Video 3 and 4:** Video depicting motion of single particles of SNAP-CD44-GFP acquired at ~60fps (3) and an example individual track (4).

**Supplementary Video 5:** Video depicting motion of single particles of SNAP-CD44TmICD-GFP acquired at ~60fps.

**Supplementary Video 6:** Video depicting motion of single particles of SNAP-CD44Tm-GFP acquired at ~60fps.

### Figure S1

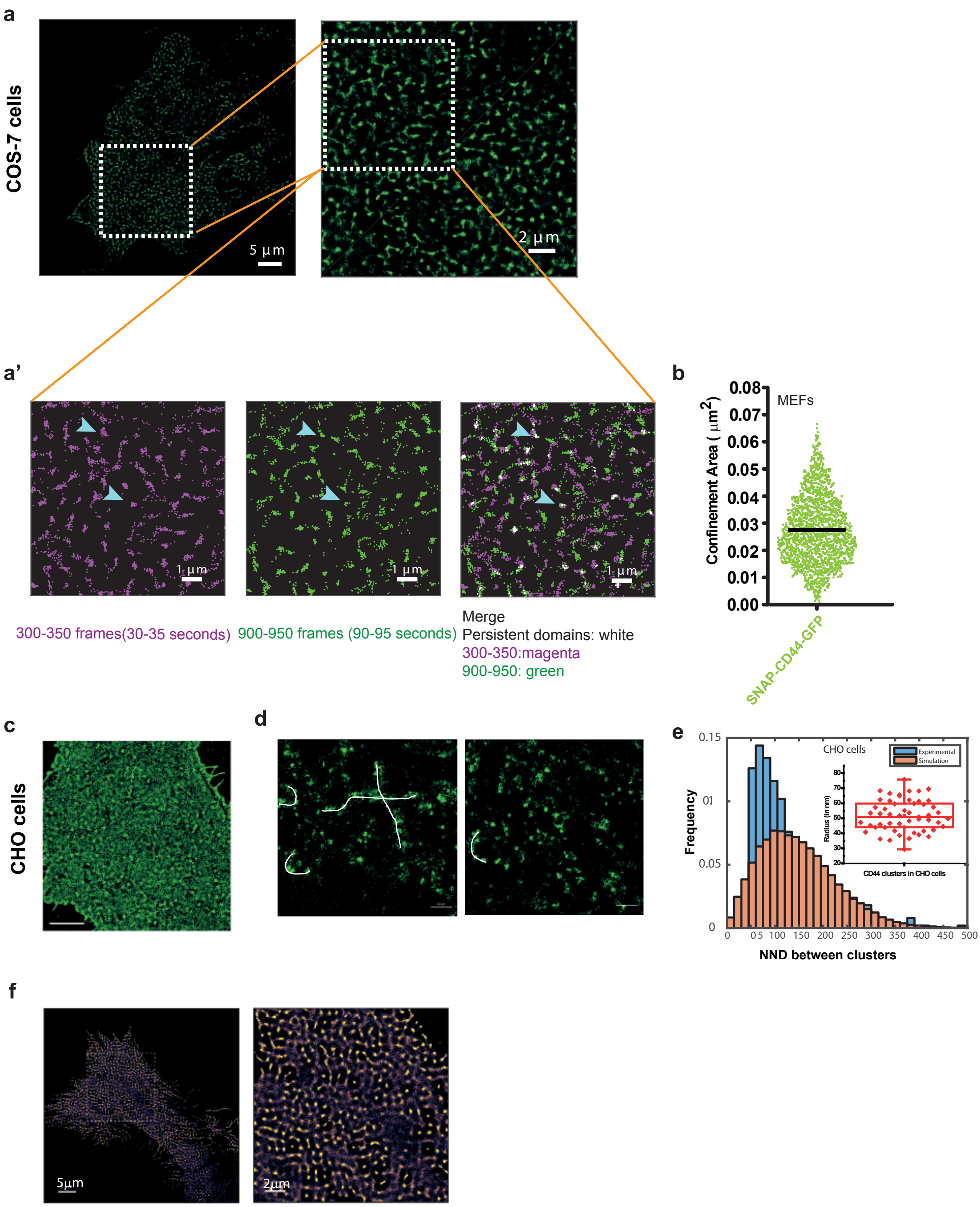
a
COS-7 cells
5 μm
2 μm
a'
1 μm
1 μm
1 μm
300-350 frames(30-35 seconds)
900-950 frames (90-95 seconds)
Merge
Persistent domains: white
300-350:magenta
900-950: green
b
MEFs
Confinement Area ( μm²)
0.08
0.07
0.06
0.05
0.04
0.03
0.02
0.01
0.00
SNAP-CD44-GFP
c
CHO cells
d
e
CHO cells
Experimental
Simulation
Frequency
0.15
01
0.05
0
0 05 100 150 200 250 300 350 400 450 500
NND between clusters
Radius (in nm)
CD44 clusters in CHO cells
f
5μm
2μm

### Figure S2

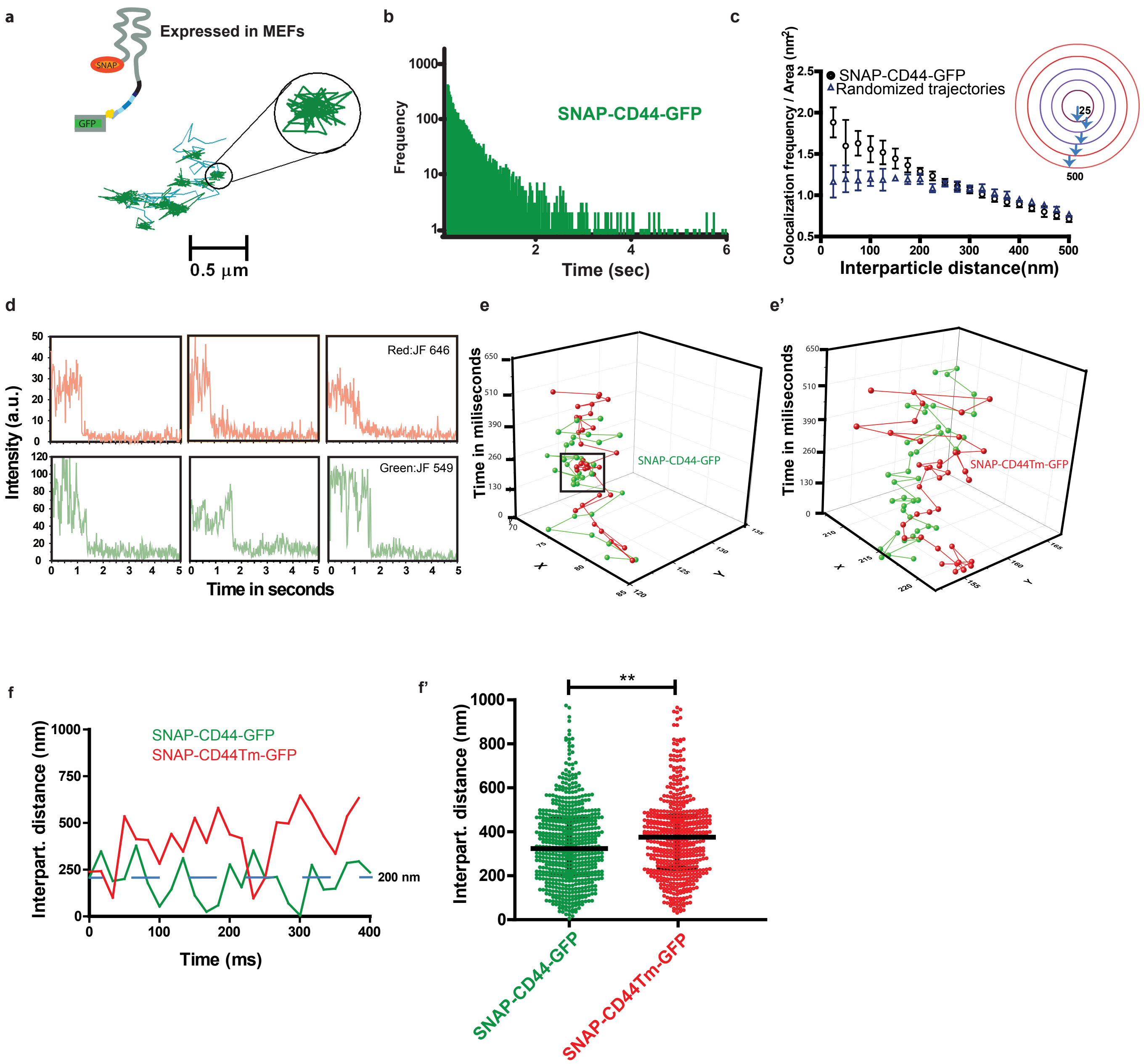

a
Expressed in MEFs
SNAP
GFP
0.5 μm
b
SNAP-CD44-GFP
Frequency
Time (sec)
c
SNAP-CD44-GFP
Randomized trajectories
Colocalization frequency / Area (nm2)
Interparticle distance(nm)
25
500
d
Intensity (a.u.)
Red:JF 646
Green:JF 549
Time in seconds
e
Time in miliseconds
SNAP-CD44-GFP
e'
Time in miliseconds
SNAP-CD44Tm-GFP
f
SNAP-CD44-GFP
SNAP-CD44Tm-GFP
Interpart. distance (nm)
200 nm
Time (ms)
f'
**
Interpart. distance (nm)
SNAP-CD44-GFP
SNAP-CD44Tm-GFP

### Figure S3

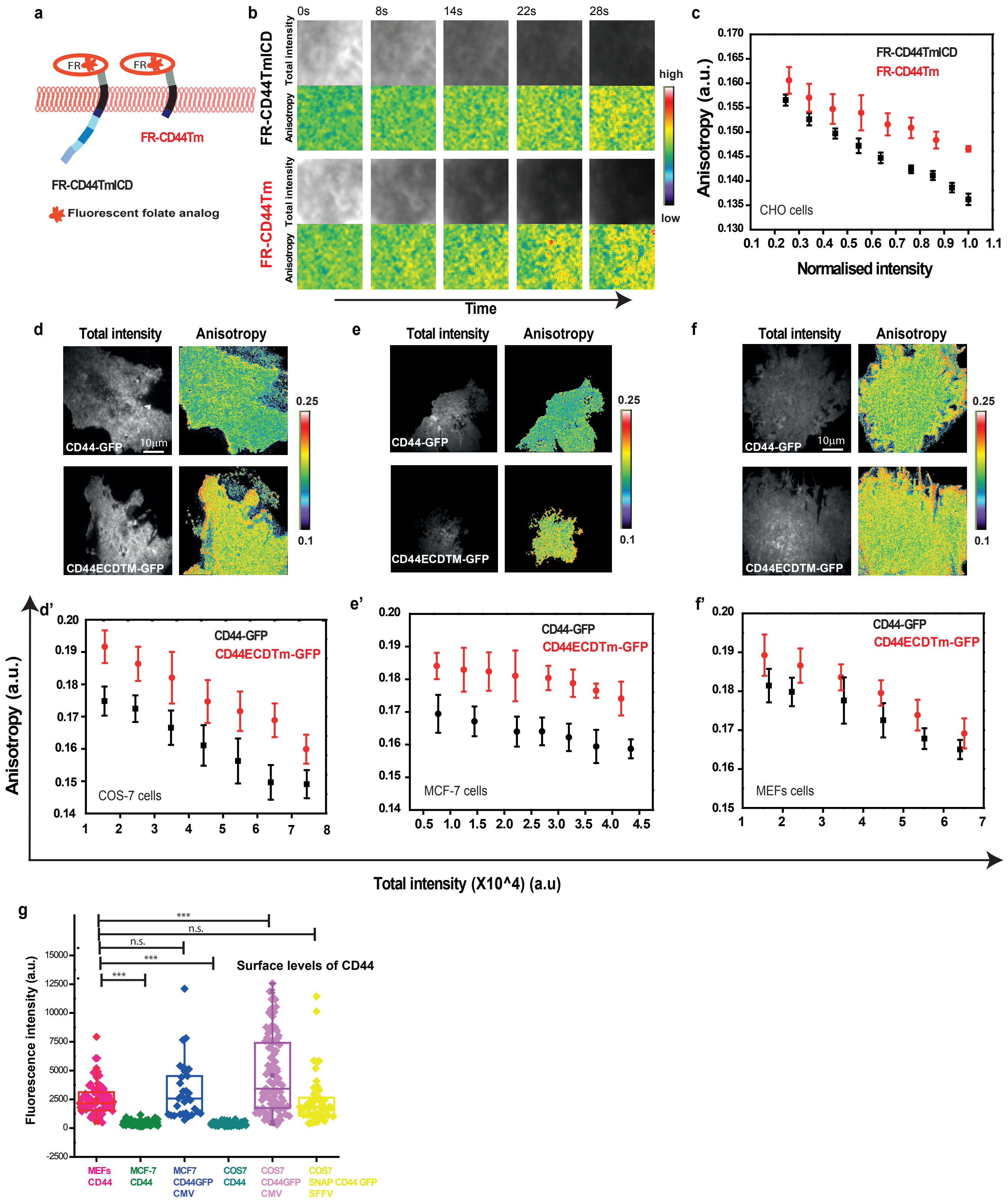
a
FR
FR
FR-CD44Tm
FR-CD44TmICD
Fluorescent folate analog
b
0s
8s
14s
22s
28s
FR-CD44TmICD
FR-CD44Tm
Total intensity
Anisotropy
high
low
Time
c
FR-CD44TmICD
FR-CD44Tm
CHO cells
Anisotropy (a.u.)
Normalised intensity
d
e
f
Total intensity
Anisotropy
CD44-GFP
CD44ECDTM-GFP
10µm
0.25
0.1
d'
e'
f'
CD44-GFP
CD44ECDTm-GFP
COS-7 cells
MCF-7 cells
MEFs cells
Anisotropy (a.u.)
Total intensity (X10^4) (a.u)
g
***
n.s.
n.s.
***
***
Surface levels of CD44
Fluorescence intensity (a.u.)
MEFs CD44
MCF-7 CD44
MCF7 CD44GFP CMV
COS7 CD44
COS7 CD44GFP CMV
COS7 SNAP CD44 GFP SFFV

### Figure S4

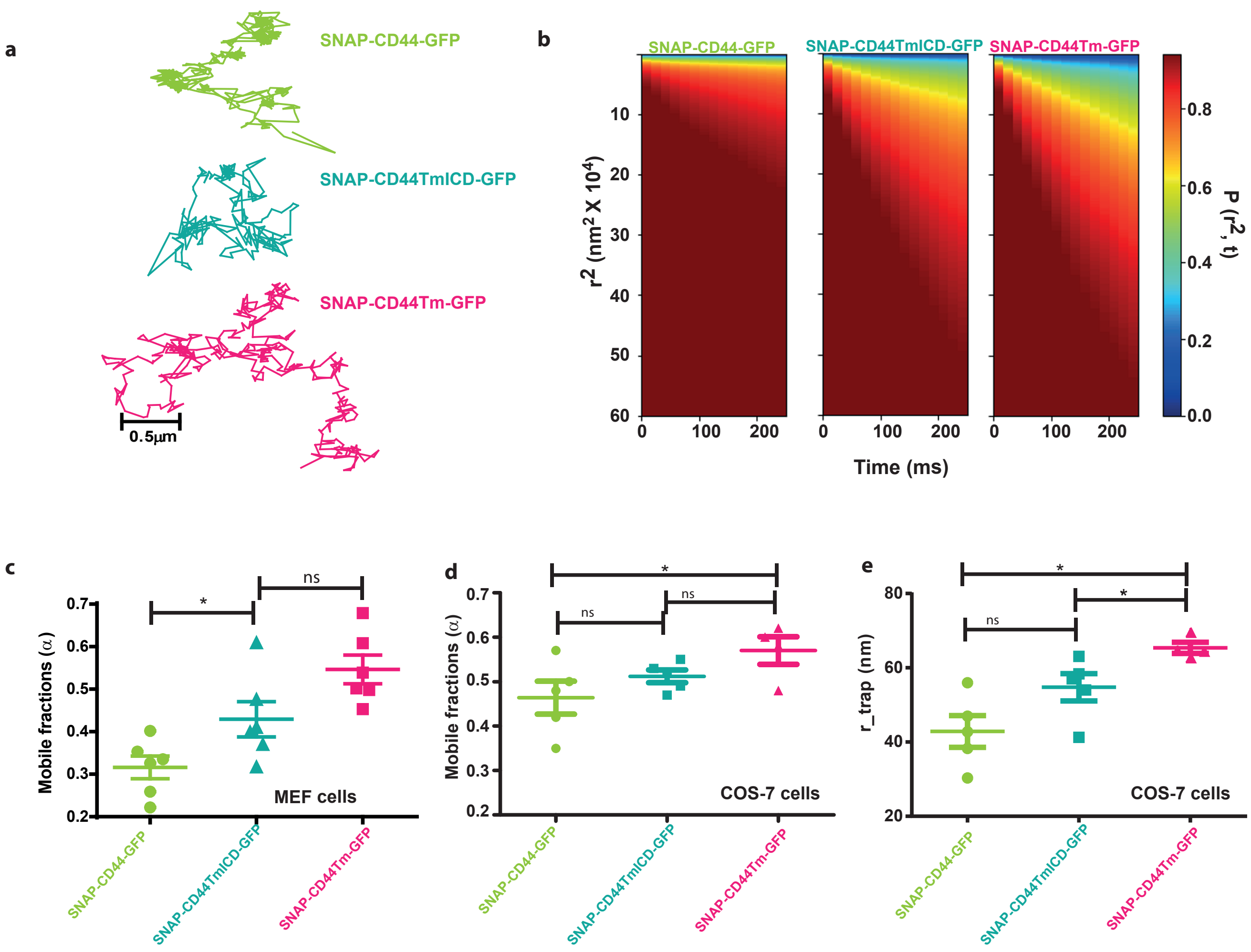

a
SNAP-CD44-GFP
SNAP-CD44TmICD-GFP
SNAP-CD44Tm-GFP
0.5µm
b
SNAP-CD44-GFP
SNAP-CD44TmICD-GFP
SNAP-CD44Tm-GFP
r2 (nm2 X 104)
Time (ms)
P (r2, t)
c
Mobile fractions (α)
MEF cells
d
Mobile fractions (α)
COS-7 cells
e
r_trap (nm)
COS-7 cells
ns
*

### Figure S5

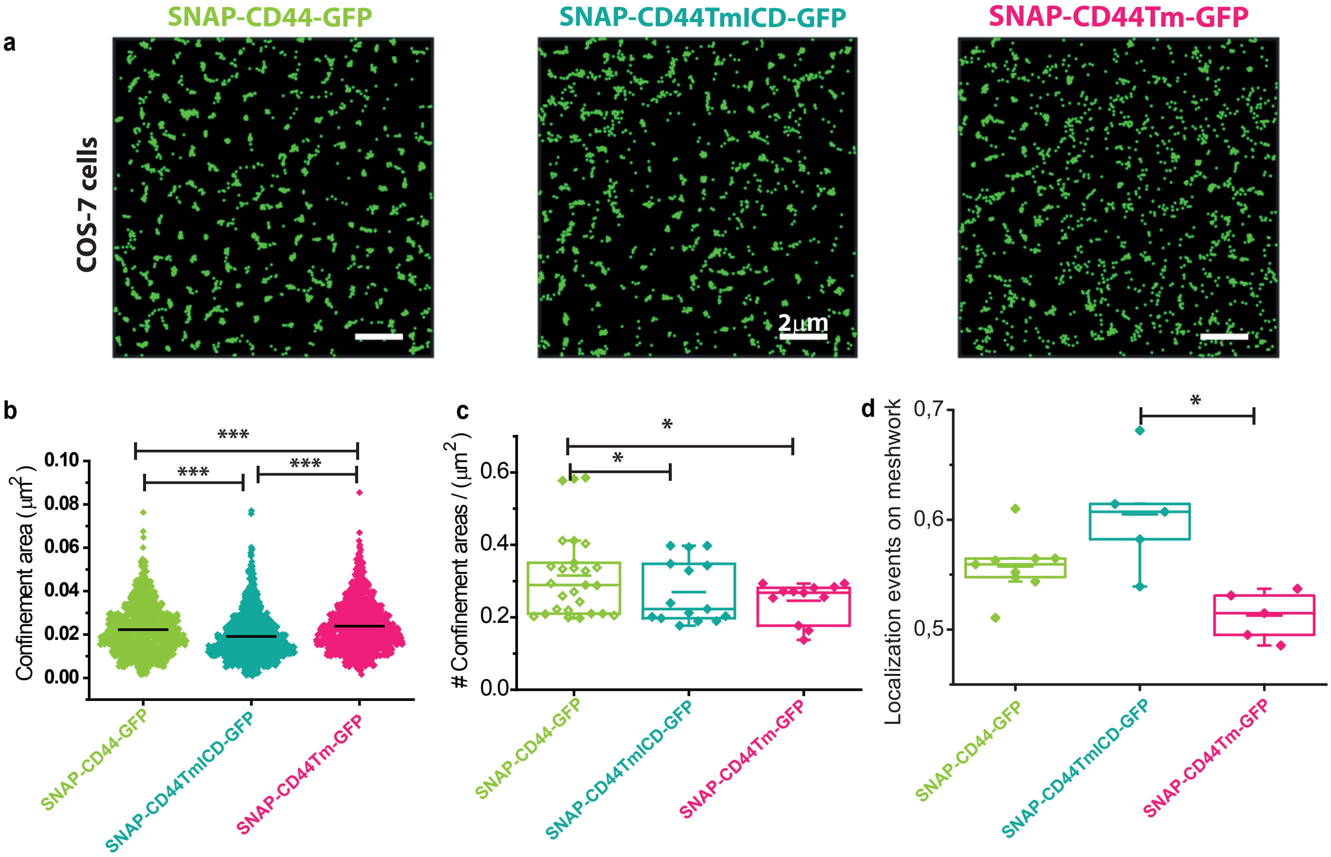

a
SNAP-CD44-GFP
SNAP-CD44TmICD-GFP
SNAP-CD44Tm-GFP
COS-7 cells
2μm
b
Confinement area (μm²)
0.10
0.08
0.06
0.04
0.02
0.00
***
***
***
SNAP-CD44-GFP
SNAP-CD44TmICD-GFP
SNAP-CD44Tm-GFP
c
### Confinement areas / (μm²)
0.6
0.4
0.2
0.0
*
*
SNAP-CD44-GFP
SNAP-CD44TmICD-GFP
SNAP-CD44Tm-GFP
d
Localization events on meshwork
0,7
0,6
0,5
*
SNAP-CD44-GFP
SNAP-CD44TmICD-GFP
SNAP-CD44Tm-GFP

### Figure S6

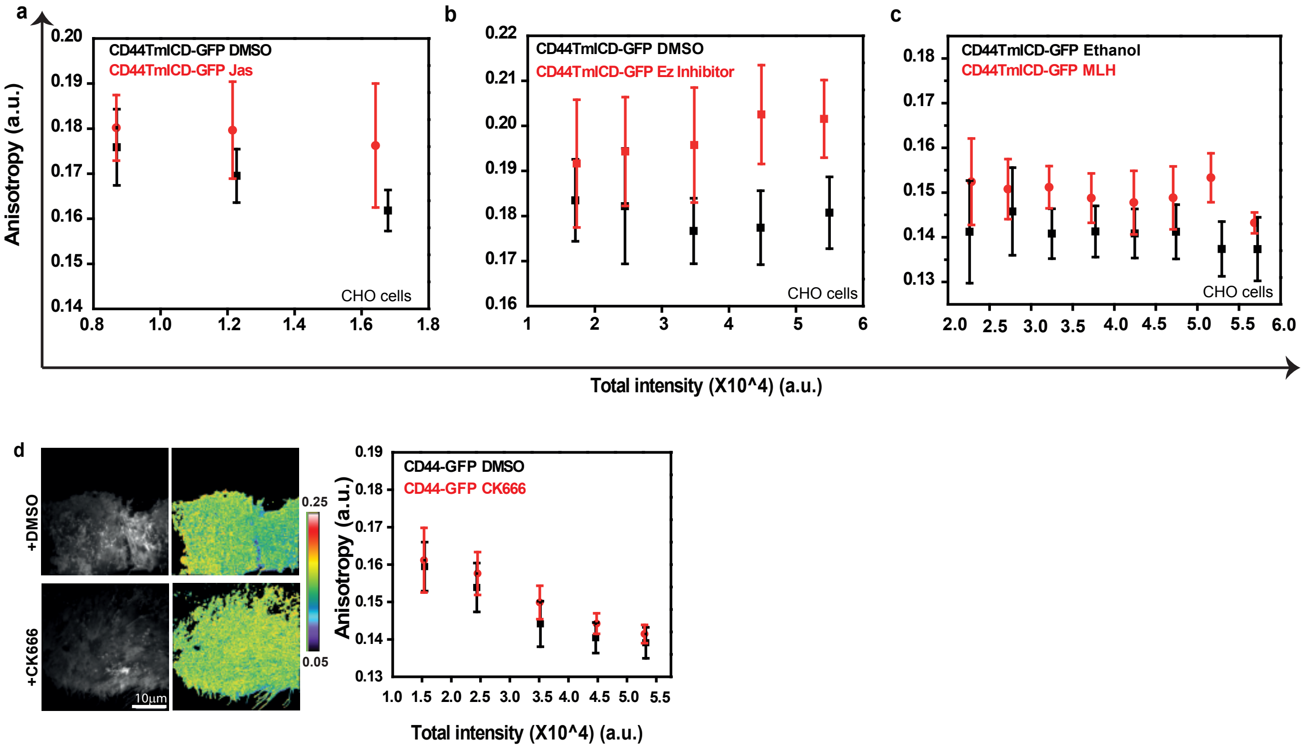
a
b
c
d
Anisotropy (a.u.)
CD44TmICD-GFP DMSO
CD44TmICD-GFP Jas
CHO cells
CD44TmICD-GFP DMSO
CD44TmICD-GFP Ez Inhibitor
CHO cells
CD44TmICD-GFP Ethanol
CD44TmICD-GFP MLH
CHO cells
Total intensity (X10^4) (a.u.)
+DMSO
+CK666
10μm
0.25
0.05
CD44-GFP DMSO
CD44-GFP CK666
Anisotropy (a.u.)
Total intensity (X10^4) (a.u.)

### Figure S7

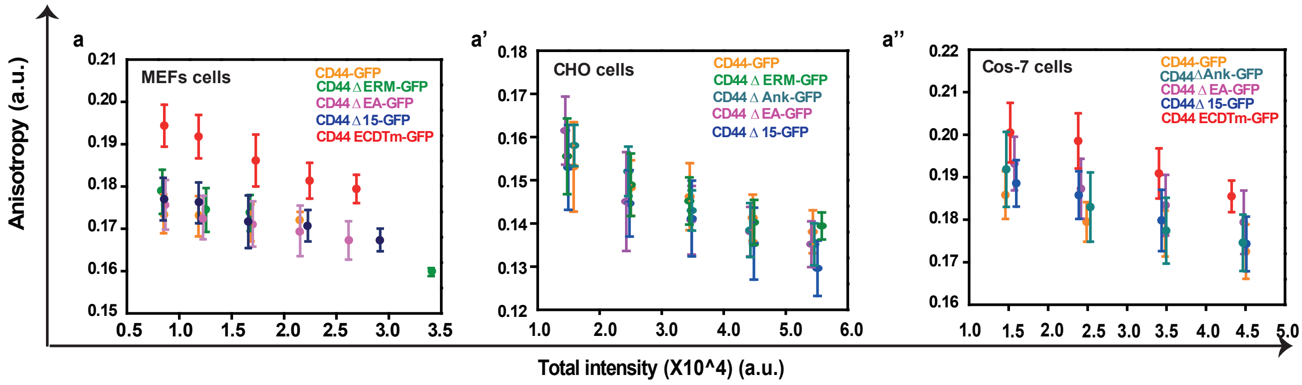

a
MEFs cells
CD44-GFP
CD44 Δ ERM-GFP
CD44 Δ EA-GFP
CD44 Δ 15-GFP
CD44 ECDTm-GFP
a'
CHO cells
CD44-GFP
CD44 Δ ERM-GFP
CD44 Δ Ank-GFP
CD44 Δ EA-GFP
CD44 Δ 15-GFP
a''
Cos-7 cells
CD44-GFP
CD44 Δ Ank-GFP
CD44 Δ EA-GFP
CD44 Δ 15-GFP
CD44 ECDTm-GFP
Anisotropy (a.u.)
Total intensity (X10^4) (a.u.)
